# Model-based and model-free valuation signals in the human brain vary markedly in their relationship to individual differences in human behavioral control

**DOI:** 10.1101/2025.09.17.676398

**Authors:** Weilun Ding, Jeffrey Cockburn, Julia Pia Simon, Amogh Johri, Scarlet J. Cho, Sarah Oh, Jamie D. Feusner, Reza Tadayonnejad, John P. O’Doherty

## Abstract

Human action selection under reinforcement is thought to rely on two distinct strategies: model-free and model-based reinforcement learning. While behavior in sequential decision-making tasks often reflects a mixture of both, the neural basis of individual differences in their expression remains unclear. To investigate this, we conducted a large-scale fMRI study with 179 participants performing a variant of the two-step task. Using both cluster-defined subgroups and computational parameter estimates, we found that the ventromedial prefrontal cortex encodes model-based and model-free value signals differently depending on individual strategy use. Model-based value signals were strongly linked to the degree of model-based behavioral reliance, whereas model-free signals appeared regardless of model-free behavioral influence. Leveraging the large sample, we found individuals lacking both model-based behavior and model-based neural signals exhibited impaired state prediction errors, suggesting a difficulty in building or updating their internal model of the environment. These findings indicate that model-free signals are ubiquitous across individuals, even in those not behaviorally relying on model-free strategies, while model-based representations appear only in those individuals utilizing such a strategy at the behavioral level, the absence of which may depend in part on underlying difficulties in forming accurate model-based predictions.

## Introduction

It has long been proposed that multiple systems underpin choice behavior (Kahneman, Frederick, et al., 2002; Loewenstein and O’Donoghue, 2004; Killcross and Blundell, 2002; Dickinson and Balleine, 2002; Rangel et al., 2008; Sloman, 1996). In many such accounts, a deliberative goal-directed system exists alongside a reflexive habitual system (Dickinson, 1985; Balleine and Dickinson, 1998; Balleine and O’Doherty, 2010; Dolan and Dayan, 2013; Daw and O’Doherty, 2014). From a computational perspective, these systems have been mapped onto two distinct forms of reinforcement-learning (RL): model-free (MF) control to account for the automatic or habitual system (Houk et al., 1994; Schultz et al., 1997, and model-based (MB) control as an account of the deliberative goal-directed system (Sutton, 2018; Daw et al., 2005; Doya et al., 2002). MF RL learns to select actions through trial and error, accumulating values for potential actions by experiencing the associated outcomes (Sutton, 1988). On the contrary, the MB system uses an internal cognitive model of the environment, which has access to the learned information of state transitions upon actions, facilitating flexible online planning (Sutton, 2018).

While human behavior has, on average, been found to reflect a mix of model-based and model-free policies (Daw et al., 2011), it is known that individuals vary considerably in the degree to which these strategies are engaged during task performance (Deserno et al., 2015; Otto et al., 2013). Some individuals exhibit mostly model-free behavior, others are more purely model-based, while yet others utilize a mixture of these two strategies (Daw et al., 2011; Wunderlich et al., 2012). Furthermore, studies have shown that there are individuals who do not utilize RL strategies at all (Jessup and O’Doherty, 2011; Colas et al., 2024; Cockburn et al., 2024).

Neural correlates of these strategies have been reported in a number of studies (Schultz et al., 1997; O’Doherty et al., 2003; McClure et al., 2003; Gläscher et al., 2010; Doll et al., 2015; D’Ardenne et al., 2008; Glimcher, 2011; O’Doherty et al., 2004; Daw et al., 2011). In particular, regions of ventromedial prefrontal cortex (vmPFC) have been found to correlate with both model-based and model-free value predictions (Hampton et al., 2006; Beierholm et al., 2011; Lee et al., 2014). However, much less is known about how individual differences in the expression of model-based and model-free behavior relate to underlying neuronal mechanisms.

It is possible that individual differences in strategy use might be reflected by differences in the degree to which a given area represents that strategy at the time of decision-making, but that the same brain areas will contribute to encoding decision-related variables irrespective of which strategy is in use. For instance, if a participant utilizes a predominantly MB strategy, perhaps the vmPFC (or another brain area) will correlate predominantly with an MB value signal, while conversely, if a participant utilizes a predominantly MF strategy, the same area will correlate instead with an MF value signal. Such a finding would indicate that MB and MF strategies are implemented within the same brain structure(s), suggesting that brain systems involved in learning and decision-making can flexibly adopt whichever strategy is being used to govern behavior. An alternative possibility is that these distinct strategies depend on distinct and dissociable brain mechanisms, such that in those individuals exhibiting MB control, unique and specific brain mechanisms are invoked to govern behavior, while in those individuals exhibiting MF control, a distinct and separate set of brain systems is engaged. A third more nuanced option is that some structures, such as the vmPFC, will flexibly encode whichever strategy is currently controlling behavior, while some other structures might exhibit strategy-specific engagement, being present only in those individuals utilizing that particular strategy to govern their behavior.

Understanding how variation in behavioral strategies across individuals relates to underlying neuronal processes has the potential to yield important insight not only into the broad question of how neuronal implementation of these computations ultimately gives rise to behavioral output, but also helps to shed light on those neural representations directly involved in guiding behavior in accordance with a particular strategy, and to separate these from those that are not directly behaviorally relevant. This research question also has the potential to help identify the neurocomputational basis of individual differences in both health and disease.

To search for distinct MB and MF signals as a function of individual differences in behavior, we implemented a version of the two-step task (Daw et al., 2011; Cockburn et al., 2024), which is structured to separate model-based and model-free behavior and its underlying neural signatures. To obtain adequate sensitivity for distinguishing between these signals across individuals, we acquired an uncharacteristically large sample (for a task-fMRI study) of participants (N=179). With the statistical sensitivity offered by such a large dataset, we therefore aimed to distinguish neural correlates of MB and MF signals in the brain and relate these to variation in behavioral strategy. We focused in particular on the vmPFC as an a priori region of interest given the extensive prior literature prominently implicating this region in both MB (or goal-directed) and MF (or habitual) valuation processes (Hampton et al., 2006; Beierholm et al., 2011; Lee et al., 2014; O’Doherty, 2011; Daw et al., 2011; Tanaka et al., 2008; Valentin et al., 2007; de Wit et al., 2009). However, we also tested for these effects across the whole brain.

## Results

### General Behaviors

Participants were instructed to perform a variant of the two-step task (Figure 1a). On each trial, participants started by choosing between two options (i.e., spaceships) at the first step, which led to one of two planets at the second stage in a probabilistic manner — one spaceship reached one planet more often than the other (common vs. rare transition in the probability of 70% vs. 30%). Then, in the second step, a transition from the planet to one of the two landing pads (also in a probabilistic manner) occurred before the participants finally observed the outcome, which was either a reward of some magnitude or no reward. Participants only had to make one choice on each trial, and the reward probability associated with choosing either of the two options fluctuated throughout the experiments (i.e., the more rewarding option would switch between the two). From the perspective of a participant, achieving a good performance depends on constantly learning the action value of the two options throughout the experiment. Representing the internal task structure when necessary (see the next paragraph, e.g., the first stage probabilistic transition from the spaceship to the planet) could also be beneficial by design when the reward is contingent upon the planet they would reach. To evaluate how well participants performed the task in general, we calculated the actual probability of obtaining a reward across all trials for each participant and obtained the population distribution of such probability of obtaining rewards. The distribution was compared to a null distribution of the probability of obtaining rewards by simulating the same number of random agents as the number of participants performing the same task. Participants’ performance was found to be significantly higher than the chance level indicated by the null distribution (*p* < 0.001, paired t-test), where the actual probability of reward from all the participants has a mean of 47.19% (*sd* = 0.393), and the null distribution has a mean of 42.75% (*sd* = 0.0278). This suggests that overall, the participants learned the reward probability of the two options well in order to obtain rewards.

**Figure 1:**
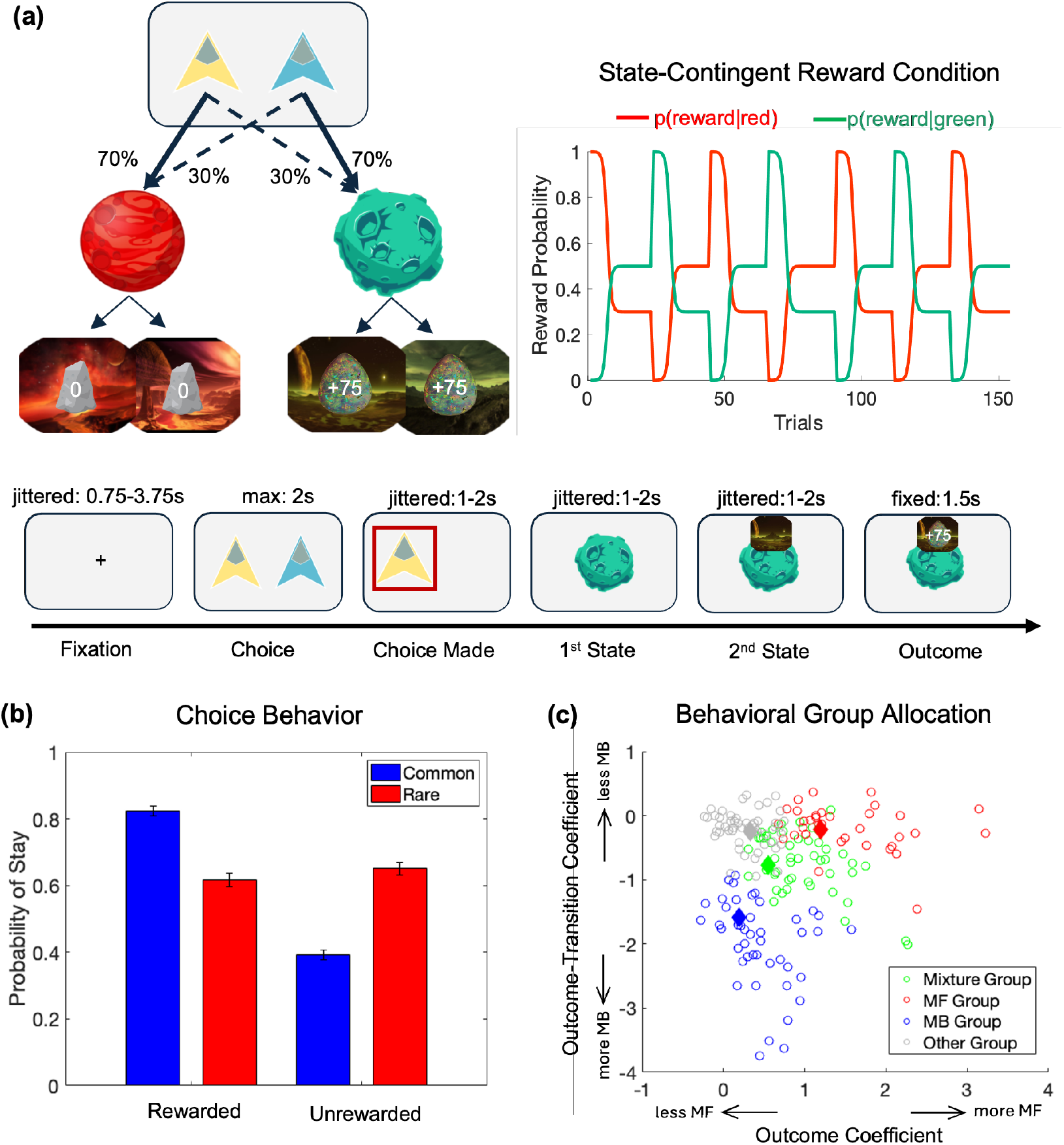
Task description and behavior. (a) Left: The two-step task structure. On each trial, participants start by choosing from two spaceships at the first step. Each spaceship has a common (70%) and rare (30%) transition to the two possible planets. Then, for the second step, the transition from the planet to the landing pads is also probabilistic, while both landing pads associated with a given planet share the same reward probability function. Right: Reward probability function throughout the task; Bottom: The timing of the trial sequence. (b) The probability of repeating the previous action as a function of the previous trial’s outcome and transition type. (c) Clustering result after allocating participants’ behavioral features to the four cluster centroids. The four diamonds denote the four centroids from the external dataset. Each empty circle denotes an individual from the fMRI sample.

Next, we aimed to determine whether participants exhibited choice behavior consistent with model-based (MB) or model-free (MF) reinforcement learning (RL) overall. In this task, as in the original two-step task (Daw et al., 2011), MF-consistent behavior can be dissociated from MB-consistent behavior by examining the choice-repeating patterns when the transition type in the preceding trial is considered. A model-free (MF) agent would choose the same spaceship as chosen on the preceding trial after receiving a reward on the preceding trial, regardless of whether a common or rare transition occurred on that trial. A model-based (MB) agent, on the other hand, would consider the nature of the state transition that occurred prior to reaching the reward on the preceding trial when making a spaceship choice: if a rare transition happened on the preceding rewarded trial, an MB agent would be less likely to repeat the spaceship choice if the reward is contingent upon the state, favoring instead the alternative spaceship associated with the common transition more likely to lead to the same state (i.e., planet) on the next trial. Prior studies have found that, on average, human behavior on this task is a mix of MB and MF strategies (Daw et al., 2011). In order to diagnose if the behaviors are consistent with MF vs. MB control, as in previous studies, we classified trials by the outcome of the previous trial (i.e., reward vs. no-reward) and the first-stage transition type (i.e., common vs. rare) in the previous trial, and we then examined the probability of choosing the same option in the current trial as in the previous trial (i.e., p(stay)) as a function of outcome and transition type in the previous trial. Qualitatively, as shown in Figure 1b, participants expressed robust sensitivity in their choice behavior to the reward, indicated by a higher probability of choosing the same option on the current trial if the chosen option was rewarded in the previous trial (averaging across preceding trials with both common and rare transitions), suggesting a model-free component in participants’ reward-maximizing strategy. In addition, participants also showed sensitivity to the transition type in the previous trial in their choice behavior. They were 1) more likely to switch their choice after a rewarded trial with a rare transition than with a common transition, and 2) more likely to stay with the choice after a non-rewarded trial with a rare transition than with a common transition, suggesting an MB component in participants’ reward-maximizing strategy. Collectively, these results support the typical observation that participants’ behavior reflects a mix of both MB and MF components on average.

To quantify the degree of reward sensitivity and transition sensitivity of the current choice, a mixed-effects logistic regression was run to test how the probability of choosing the same option is influenced by the previous outcome, previous transition type, and the interaction between these variables (see Methods for details). Consistent with the qualitative behavioral results, MF-consistent behavior was observed, in that participants are more likely to repeat their choice if that choice was rewarded in the previous trial, reflected by an MF-consistent main effect of the previous outcome (*β* = 0.721, *SE* = 0.0527, *T* = 13.695, *p* < 0.001). MB-consistent behavior was also found in that participants are more likely to switch to the other spaceship when they experience a rare transition towards the reward, reflected by a significant effect of the interaction between the previous outcome and the previous transition type (*β* = −0.751, *SE* = 0.0673, *T* = −11.157, *p* < 0.001). In sum, the participants showed a mixture of MF-consistent and MB-consistent behaviors, replicating the typical behavioral patterns observed in the two-step task (Daw et al., 2011).

### Computational Variables and Behavioral Clusters

Though participants expressed a mixture of MF and MB strategies at the population level, it is also possible that participants vary substantially in their behavioral strategies. In a companion study using the same task in a larger behavioral sample (N=678; Cockburn et al., 2024), we found evidence for distinct behavioral clusters that reflect individual differences in the use of underlying strategies. In that study, we reported four distinct clusters corresponding to distinct behavioral profiles: a predominantly MB group, a predominantly MF group, those exhibiting a mixture of the two strategies, and a fourth group that did not appear to rely on RL-like mechanisms at all. Here, we utilized the same clustering features as used in that previous study. The features in question include statistics about stay-switch choices and reaction times as a function of the previous trial’s event (switch vs stay decisions). We relied on the same four cluster centroids obtained from that previous study to assign the participants in the current sample to the corresponding four groups (see “Behavioral Clustering” in the Methods section for more details). As a result we have the composition as follows: 1) a Mixture Group (N=48), 2) an MF Group(N=34), 3) an MB Group(N=44), and 4) an Other Group (N=53). The composition of the four groups is shown in Figure 1c as projected onto the dimensions of the outcome and outcome-transition coefficients.

When evaluating the performance of the obtained four behavioral groups, we found that the Mixture Group (49.55%, *sd* = 0.0291), MF Group (47.27%, *sd* = 0.0353), and MB Group (47.94%, *sd* = 0.0381), were significantly better than random agents whereas the Other Group (44.37%, *sd* = 0.0342) showed a slightly better-than-chance performance, suggesting relatively blunted reward-driven behaviors (Figure S1a).When plotting each group’s probability of stay choice as a function of the previous trial’s outcome and transition type, as shown in Figure 2, we observed that the MF group repeats the previous rewarded choice and switches away from the unrewarded choice irrespective of whether the previous trial’s transition is common or rare; in contrast, the MB group repeats the rewarded choice after a common transition or an unrewarded choice after a rare transition in the previous trial, and switches away from the same option after an unrewarded common-transition trial or a rewarded rare-transition trial. The Mixture group expresses stay or switch choice patterns at the intermediate level between the MF and the MB group, whereas the Other group shows choice patterns that are relatively insensitive to the outcome, regardless of the transition type in the previous trial. Consistently, when examining the random effects of the outcome coefficient across the groups, we found a significant linear trend indicating a stronger outcome effect going from MB to MF groups (*MB < Mixture < MF, β* = 0.781, *SE* = 0.0825, *T* = 9.469, *p* < 0.001), while the Other group shows a relatively weak outcome effect compared to the RL groups (*Other < MB*, two-sample t-test, *p* = 0.0201, *Other < Mixture*, two-sample t-test, *p* < 0.001; *Other < MF*, two-sample t-test, *p* < 0.001). Similarly, for the random effects of outcome-transition interaction, the effect became stronger when moving from MF to MB groups (*MF < Mixture < MB, β* = −0.690, *SE* = 0.0476, *T* =− 14.492, *p* < 0.001), while the Other group shows a relatively weak outcome-transition effect compared to the RL groups, comparable to that of the MF group (*Other* ≈ *MF*, two-sample t-test, *p* = 0.8199; *Other < Mixture*, two-sample t-test, *p* < 0.001; *Other < MB*, two-sample t-test, *p* < 0.001).

**Figure 2:**
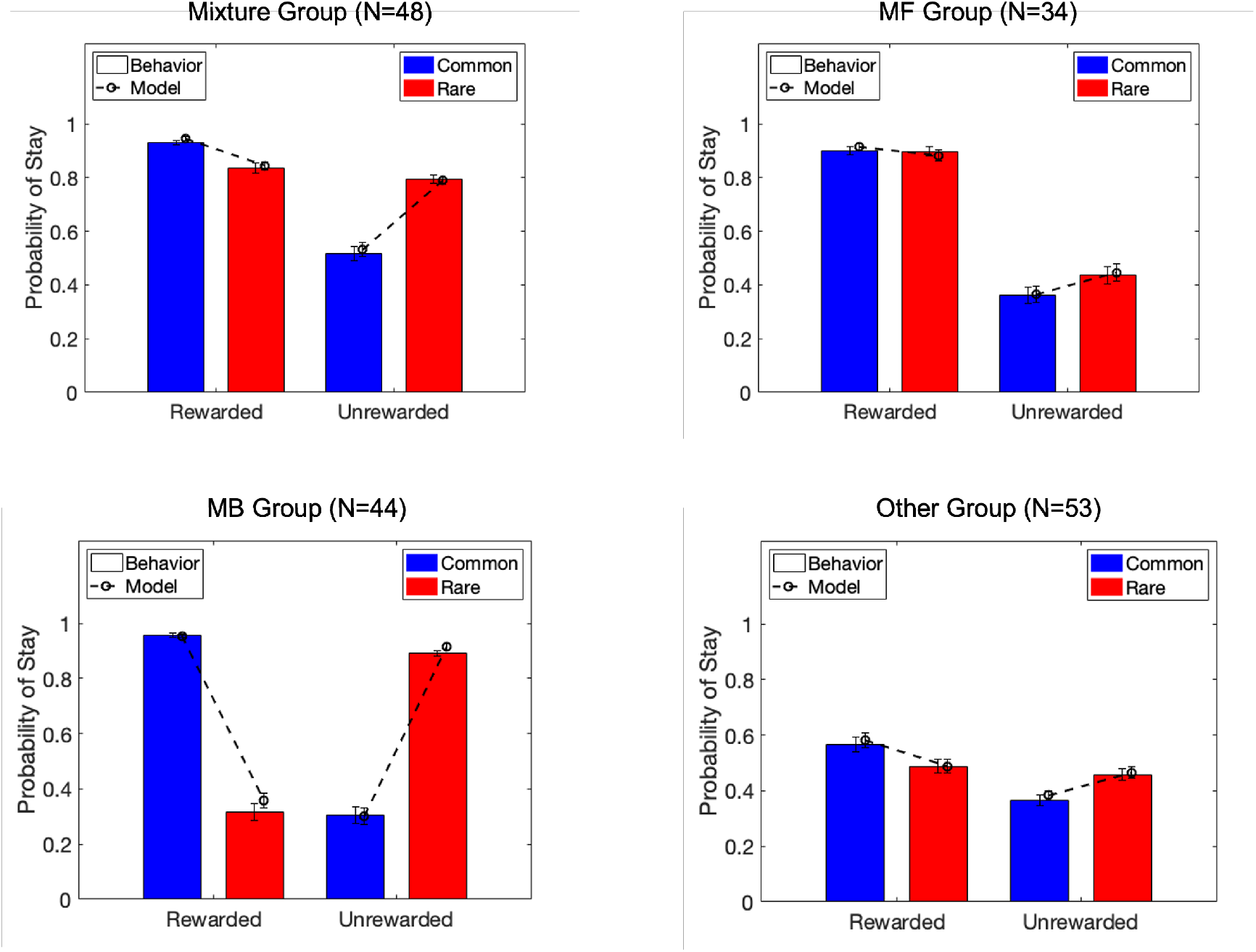
Behaviors and model predictions for individual behavioral groups. The stayswitch choice behavior for each behavioral group using group memberships shown in Figure 1c. The bars indicate the participants’ actual behavior, and dashed lines illustrate the predicted behavior from the fitted computational model. The error bars reflect across-subject SEMs.

After confirming the validity of the Mixture, MB, and MF and Other group assignments, we next sought to characterize the key MF and MB computations within each corresponding system that facilitate the expression of MF and MB behavioral signatures. For this purpose, we fit a hybrid mixture reinforcement learning model (which we called the Arbitration Mixture model) with separate MF and MB modules learning over both slow and fast timescales (Iigaya et al., 2019) and with independent condition-specific weights assigned to the MF and MB systems (Cockburn et al., 2024; see Methods for details). The model’s posterior predictions align with the participants’ actual behavior quite well (Figure 2), which indicates good model performance, thereby validating the use of learning and decision variables provided by this model to further probe their neural substrates. Systematic model comparison between this model and alternative models was performed and detailed extensively in the same previous study using a large behavior-only dataset as described earlier (Cockburn et al., 2024). In that study, the arbitration mixture model fit the data better than the alternatives. We repeated the model comparison process here using the AIC metric (Schwarz, 1978) on the current dataset, and similarly found that the arbitration mixture model explains the behavioral data the best (see Table 1).

**Table 1.** Model AIC scores in explaining the 2-step task behavior. The arbitration mixture model had significantly lower AIC scores than the other models, F(3,712) = 7.84, *p* < 0.001; paired-sample t-tests of Arbitration Mixture vs. each of the other models were significant at *p* < 0.001)

| Model | AIC |
| --- | --- |
| Arbitration Mixture | 243.0035 |
| Fixed-weight Mixture | 245.4353 |
| Model-Free | 274.3900 |
| Model-Based | 266.3880 |

Within this mixture model, the Q-learning algorithm for the MF system is used to learn the magnitude of the outcome and the expected state values of each intermediate stage. On each trial, as there are state transitions towards three different stages (i.e., planet, landing pad, and outcome), we modeled three distinct MF reward prediction errors (RPE) occurring within a trial, approximating what would be expected from a temporal difference learning rule (Sutton, 2018). Specifically, the MF RPE is assumed to be computing the difference between the expected state value and the currently encountered state value, with the state value at the outcome stage corresponding to the received reward magnitude. As for the MB system, two sets of state prediction errors (SPE) are computed at the two transition (i.e., spaceship-to-planet, and planet-to-pad) with experiencing a low-probability transition would signal a large SPE. The experienced SPEs are then used to compute the MB state value at the first and second stages, respectively, through computing the expected value by considering the probability of going into each state. The terminal state’s value (pad value) is learned through the MB RPE computed in the same way as for the MF RPE experienced during the outcome stage with the difference being that the MB RPE considers binary outcome yet the MF RPE considers the magnitude outcome (see “Computational Models” in the Methods section for details).

The average trial-wise MF weight across all participants is *wMF* = 0.5322 (*SE* = 0.0173), and the MF weights for the 4 sub-groups correspond well with their cluster labels - Mixture group: *wMF* = 0.6436 (*SE* = 0.0259); MF group: *wMF* = 0.7332 (*SE* = 0.0228); MB group: *wMF* = 0.2684 (*SE* = 0.0210); Other group: *wMF* = 0.5215 (*SE* = 0.0239) (see Figure S1b for the MF weight distributions for four groups; see Tables S1-S6 for details of model parameters).

### Neural correlates of model-based and model-free decision value in general

To address which RL computations are carried out in the brain to give rise to the different behavioral strategies, we performed an analysis of the fMRI data in which we included decision value variables from both MF and MB systems to test for their neural correlates (see Methods: Computational models for details of the decision value variables). The decision value we examined is defined as the value of the chosen option minus the value of the rejected option (the value here contains both the Q-value from the respective RL model and another decision component). We entered both the chosen and the rejected option value from the MF and MB systems simultaneously as parametric modulators for the stick function at the time of stimulus onset (i.e., spaceship onset). We then used the contrast of chosen value minus rejected value to examine the effects of decision value from the MF and MB systems in the brain.

First, we tested for the effect of MB and MF decision value across participants (pooling across all participants in the study who had behavior consistent with RL strategies, i.e., excluding those in the non-RL Other group). We found a significant effect of MB decision value in the vmPFC (*p* = 0.002, cluster-level FWE-corrected, Figure 3a left; similar effects were found even if also including participants in the non-RL other group, *p* = 0.001, cluster-level FWE-corrected). Outside of the vmPFC, we also found significant MB decision value effects in the mid-cingulate cortex (MCC), precuneus cortex (an inter-region cluster including the occipital cortex), and right amygdala (all *p* < 0.001, cluster-level FWE-corrected, Figure 3a; see Table S7 for a full list of brain regions showing significant MB decision value effects at *p* < 0.05, whole brain FWE-corrected). After identifying the brain regions showing a significant MB decision value effect, we then examined where in the brain the MB decision value effect was correlated with the behavioral tendency of showing MB (or MF) behavior with a parametric variable, *wMF*, from the computational model in the second-level GLM analysis. We observed that, in the vmPFC, participants with a higher behavioral MB tendency (i.e., smaller *wMF*) showed stronger MB decision value effects (*p* < 0.001, cluster-level FWE-corrected, Figure 3b). Beyond the vmPFC, similar patterns were observed in the precuneus (and posterior cingulate) cortex (*p* < 0.001, cluster-level FWE-corrected, Figure 3b; see Table S9 for details).

**Figure 3:**
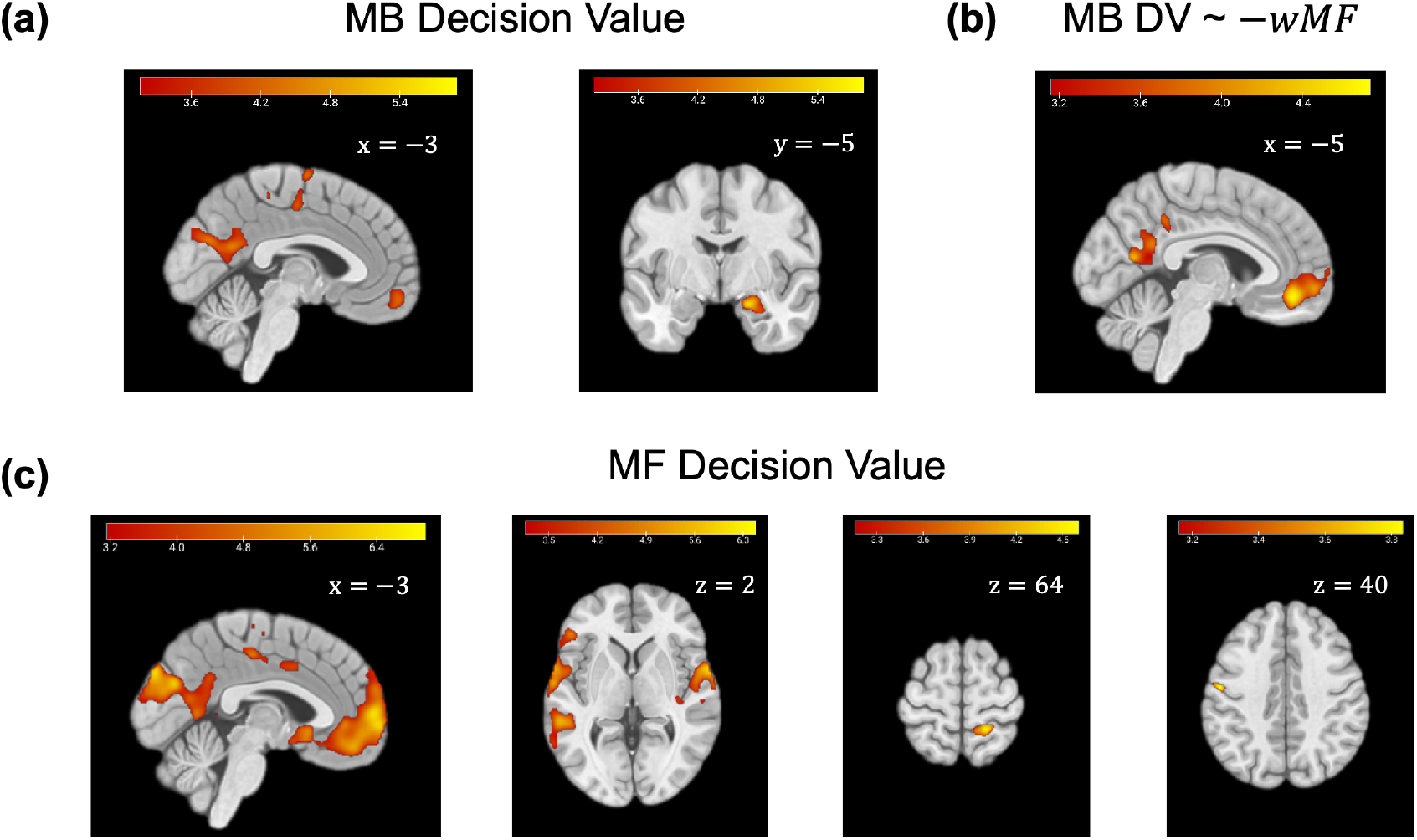
Encoding of MB and MF decision value at the group level, and across individuals. All brain images show the T-maps with a cluster-forming threshold of *p* < 0.001, and cluster-level FWE-corrected *p* < 0.05. (a) Clusters with activities correlating with the MB decision value across participants using RL strategies; left: ventromedial prefrontal cortex (vmPFC), supplementary motor (and mid-cingulate) cortex, and precuneus (and occipital) cortex; right: right amygdala. (b): Clusters in vmPFC, and precuneus (and posterior cingulate) cortex whereby the MB decision value correlates with the degree of MB control (-wMF) across participants. (c): Clusters with activities correlating with the MF decision value across participants using RL strategies; left: ventromedial prefrontal cortex (vmPFC) including subcallosal cortex, mid-cingulate cortex, precuneus cortex, and occipital cortex; mid-left: left central opercular cortex, left ventrolateral prefrontal cortex, and right central opercular cortex; mid-right: right postcentral gyrus; right: left precentral gyrus.

We then implemented the same analyses, but this time examining MF value representations. When pooling over all RL participants in the study (excluding participants in the non-RL other group), we found a significant effect of MF decision value (i.e., chosen value minus rejected value) in the vmPFC (including the subcallosal cortex, *p* < 0.001, cluster-level FWE-corrected, Figure 3c left; similar results were observed even if including participants in the Other group as well, *p* < 0.001, cluster-level FWE-corrected). In regions outside of the vmPFC, the mid-cingulate cortex(MCC), the precuneus cortex, and the occipital cortex were also found to show significant MF decision value effects (an inter-region connected cluster, *p* < 0.001, cluster-level FWE-corrected, Figure 3c left). MF decision value effects were also observed in the left central opercular cortex and left ventrolateral prefrontal cortex (an inter-region connected cluster *p* < 0.001, cluster-level FWE-corrected, Figure 3c mid-left), in the right central opercular cortex (*p* < 0.001, cluster-level FWE-corrected, Figure 3c mid-left) as well as in the right postcentral gyrus (*p* = 0.002, cluster-level FWE-corrected, Figure 3c mid-right) and the left precentral gyrus (*p* = 0.049, cluster-level FWE-corrected, Figure 3c right; see Table S8 for a full list of brain regions showing significant MF decision value effect at cluster-level FWE-corrected *p* < 0.05). When testing for correlations between the behavioral MF tendency (*wMF*) and the MF decision value effects, no significant clusters in the brain were observed surviving cluster-level FWE-correction (*p* < 0.05 with a cluster-forming threshold of *p* < 0.001), indicating no significant relationship between MF decision value effects and the degree to which MF strategies are present in behavior across participants.

For a complete illustration of the overall decision value effect presented here, we also included the neural effects of the sub-component of decision values - the chosen and rejected values of the MB and MF systems in the supplements (Figure S3).

### Sub-group differences in neural correlates of model-based and model-free decision value

In addition to examining the overall MB and MF decision value signal across all individuals who show strong RL signature, we further examined correlates of MB and MF decision value signals in each behavioral cluster group at the whole-brain level. We found robust correlates of an MB value signal in vmPFC (*p* < 0.001, cluster-level FWE-corrected) in the MB group (Figure 4a; see Table S10 for all brain regions showing significant MB value signals in the MB group). We observed weaker correlations with MB value in the other three groups, with modest effects of MB value in the Mixture group (at *p* < 0.01, uncorrected) and no clusters entirely located in the grey matter of vmPFC surviving at *p* < 0.01 (uncorrected) in the MF and Other groups (Figure 4a; see Table S11 for all brain regions showing significant MB value signals in the Mixture group).

**Figure 4:**
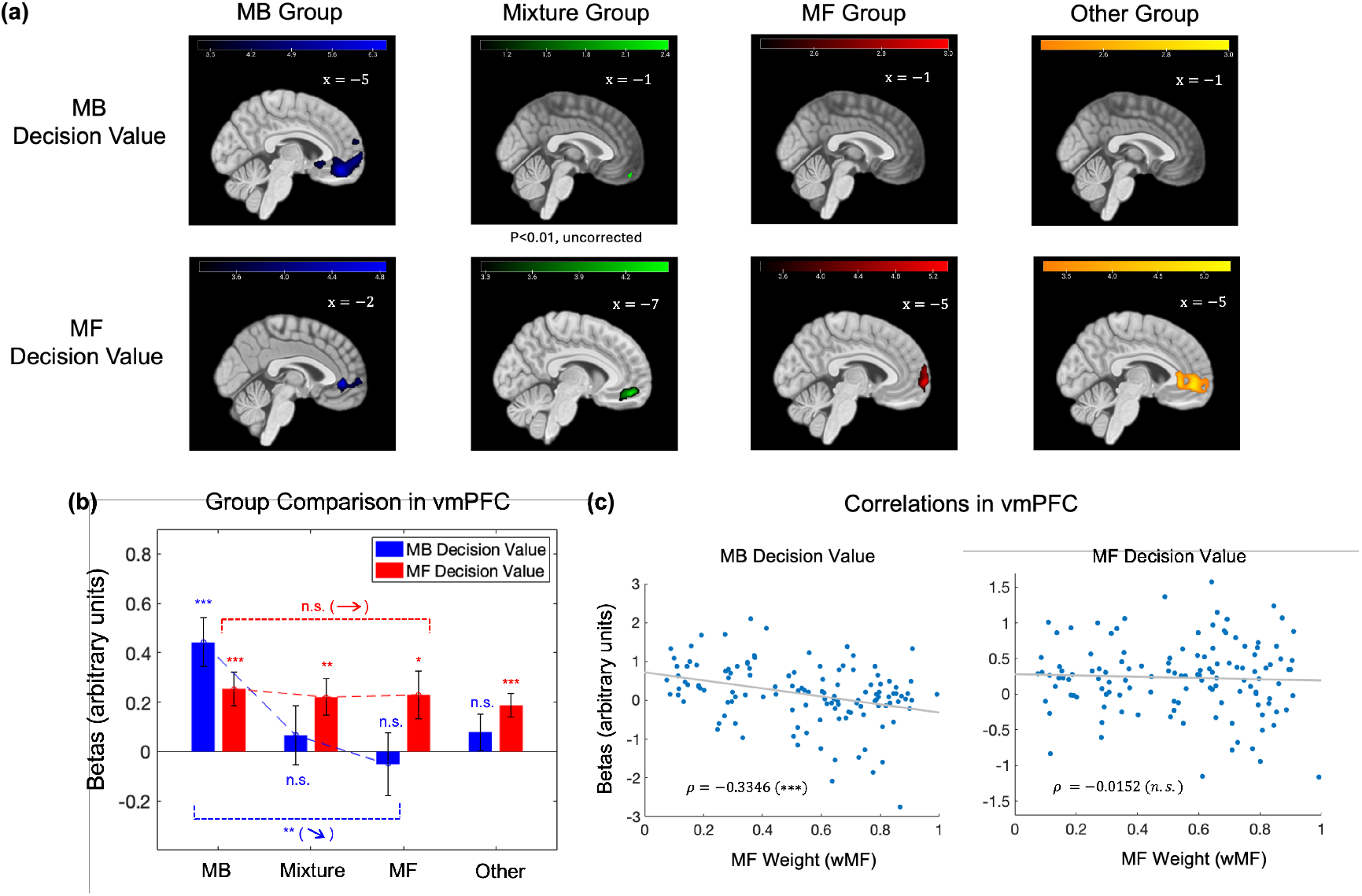
Encoding of MB and MF decision value at the sub-group level, and across individuals. All brain images show the T-maps with a cluster-forming threshold of *p* < 0.001, and cluster-level FWE-corrected *p* < 0.05 unless otherwise indicated. (a) Clusters in vmPFC/mPFC correlating with MB and MF decision value within each behavioral sub-group; top: clusters in vmPFC/mPFC correlating with MB decision value show a strong effect in MB Group, relatively modest effect in Mixture Group but weak/null effect in the MF or Other group (Mixture, MF and Other Groups’ brain maps are shown at a cluster-forming threshold of *p* < 0.01, uncorrected); bottom: clusters in vmPFC/mPFC correlating with MF decision value show strong effects in all four groups (including the Other Group). (b): Parameter estimates of MB decision value (blue) and MF decision value (red) in the vmPFC ROI from all behavioral groups (statistics are from the one-sample t-test on whether the betas in vmPFC are significantly larger than zero for each group), along with the ordinal group effect of the MB decision value (blue dash line, significantly decreasing from MB to Mixture to MF groups) and MF decision value (red dash line, non-significant change across MB, Mixture and MF groups) run through a regression model across the three RL groups. (c): Spearman correlations between the beta coefficients of MB decision value (left) and MF decision value (right) in the vmPFC ROI and the weight of the MF system from the computational model. To denote statistical significance: ***: *p* < 0.001, **: *p* < 0.01, *: *p* < 0.05, n.s. : *p >* 0.05.

Next, we tested for MF decision value signals in each behavioral cluster group separately. In contrast to the neural representation of MB decision value, MF decision value was represented significantly in all three RL-related cluster groups and even in the non-RL (Other) group in vmPFC (cluster-level FWE-corrected, MF Group: *p* < 0.001; Mixture Group: *p* < 0.001; MB Group: *p* < 0.001; Other Group: *p* < 0.001; Figure 4a; see Tables S12-S15 for all brain regions showing significant MF value signals in each sub-group), in the vmPFC, implying that MF decision value is also represented in the groups whose behaviors align more with a pattern of MB control or even in the group using a predominantly non-RL (or a blunted RL) strategy.

To perform a more detailed analysis on the vmPFC, we defined an independent ROI utilizing a mask derived from an independent meta-analysis involving more than 200 fMRI studies examining the neural correlates of value (Bartra et al., 2013), and performed a direct statistical comparison across groups within this ROI. To estimate the overall signal correlations in the vmPFC ROI and to compare the representation of decision values of MF and MB systems from group to group, we ran a regression model on the 1st-level beta estimates in the vmPFC (ROI) with an intercept and an ordinal group variable on MB decision value (see Methods). We found a significant overall positive effect of MB decision value indicated by a significant intercept (*β* = 0.651, *SE* = 0.178, *T* = 3.66, *p* < 0.001). However, we also found that MB decision value was represented in vmPFC differently across the three groups, indicated by a significant ordinal group effect (*β* = −0.253, *SE* = 0.0857, *T* = −2.95, *p* = 0.0038, Figure 4b). In particular, the representation of MB decision value in vmPFC was most strongly represented in the MB group, but as the MB contribution to the behavior shrank, the neural representation of MB decision value in vmPFC waned.

We next used the same vmPFC ROI, as in the MB value analysis, for the analysis of the MF value signals. Mirroring the results of the whole-brain analysis, we found that MF decision value was represented significantly in all three groups using RL strategies — in the MB group (*p* < 0.001, one-sample t-test), Mixture group (*p* = 0.0040, one-sample t-test), and MF group (*p* = 0.0229, one-sample t-test), as well as in the non-RL Other group (*p* < 0.001, one-sample t-test). We ran a regression model on the 1st-level beta estimates of MF decision values in the vmPFC (ROI) with an intercept and an ordinal group variable (see Methods). Consistent with the whole-brain fMRI analysis, we found, in the vmPFC ROI, significant overall positive correlations for MF decision value indicated by a significant intercept (*β* = 0.260, *SE* = 0.118, *T* = 2.20, *p* = 0.0295). We then tested for an ordinal effect of MF decision value, which revealed no significant evidence for an ordinal group effect in the data (*β* = −0.0131, *SE* = 0.0569, *T* = −0.231, *p* = 0.8181, Figure 4b). This further demonstrates that the representation of MF decision value in vmPFC across the three RL groups is comparable regardless of the extent to which model-free behaviors are expressed.

To further test whether variation in MB/MF behavioral control is related to variation in the representation of MB and MF decision value signals in the vmPFC without relying on categorical group assignments, we computed Spearman correlations between each participant’s model-weighting parameter (*wMF*) derived from the behavioral computational model fits (capturing how MB or MF an individual is) and their MB and MF value signals in the same vmPFC ROI. Across the whole sample, we found a significant correlation between the representation strength of MB decision value in vmPFC and each individual’s MB weight, such that the more MB they are, the stronger the effect of MB value in vmPFC, and the less MB they are, the weaker the effect of MB value in vmPFC (Spearman’s *ρ* = −0.3346, *p* < 0.001, Figure 4c left). In contrast, we found that when the individual’s behavior went from MB to MF (with increasing *wMF*), the representation of MF decision value did not shift significantly (Spearman’s *ρ* = −0.0152, *p* = 0.8658, Figure 4c right). In other words, MF value signals are persistently represented in the vmPFC across individuals independently of the degree to which MF signals actively contribute to behavior. For instance, even in people exhibiting a strong MB strategy in their behavior, activity in the vmPFC nevertheless was found to be correlated with the MF strategy.

### Comparison of model-free and model-based decision value across MB and MF groups

We also conducted a two-sample t-test to directly compare the MB group and MF groups in terms of whether they represent MF decision value and MB decision value differently. These statistical comparisons confirm that MB decision value is more strongly represented in the MB group than in the MF group in vmPFC (*p* < 0.001, cluster-level FWE-corrected, see Table S16 for details and other brain regions involved), yet there is no significant difference in vmPFC activity correlating with MF decision value between the MB and MF groups. Furthermore, we looked into the independent vmPFC ROI (Bartra et al., 2013) and extracted beta estimates of decision values there on which we performed a two-way ANOVA to examine the effects of behavioral group (i.e., MB or MF group) and value type (i.e., MB or MF decision value), and we found a significant group and value-type interaction effect (*F* (1, 152) = 5.74, *p* = 0.0178). Based upon the ANOVA results, we further found that in the vmPFC ROI, consistent with the whole-brain statistical parametric comparison, the MB group represents MB decision value significantly stronger than the MF group does (two-sample t-test, *p* = 0.0027), and there is no significant difference in terms of MF decision value representation between the two groups (two-sample t-test, *p* = 0.8328).

We would like to note that, to take into account the potential confounding effect of age on our findings, in an additional analysis we reran the 2nd level fMRI analyses while including age as a covariate for all relevant analyses on the MB and MF decision values. The results presented in the previous and current sections still hold up at the relevant statistical thresholds.

### State prediction error signals and the relevance with behavioral control

Next, we tested for BOLD correlates of prediction error signals, which are key signals suggested to be involved in learning the representations required for implementing MF and MB strategies (Sutton, 1988; Sutton, 2018; Gläscher et al., 2010; Möhring and Gläscher, 2023). In particular, two signals have been reported in the literature as being involved in MB and MF learning, respectively: the state-prediction error (SPE) and the model-free reward prediction error (RPE). The SPE is suggested to mediate learning of the state-action-state transitions underpinning the internal model used by the MB system to compute value signals (Gläscher et al., 2010; Lee et al., 2014) and the MF RPE is suggested to support learning of MF value signals (Sutton, 1988; Montague et al., 1996; Schultz et al., 1997; Waelti et al., 2001; O’Doherty et al., 2003). Another RPE signal that has been reported in the literature is the MB RPE (Daw et al., 2011). While the MB RPE is not strictly necessary for learning within an MB system (though it can be so deployed under some implementations), it can serve as an indicator of whether or not a planning strategy has been successful or not in terms of leading the agent to a rewarding outcome. In our fMRI analysis, we included the SPE signals at both first-stage adn second-stage transitions and also included both the MF RPE and MB RPE signals as parametric regressors, pooling across all timepoints within a trial at which RPE signals were modeled.

As for SPE signals, we found evidence at the whole-group level (all groups using RL strategies) for these signals in both dorsolateral prefrontal cortex (bilateral dlPFC, *p* < 0.001, cluster-level FWE-corrected, Figure 5a top left) and intraparietal sulcus (bilateral IPS, *p* < 0.001, cluster-level FWE-corrected, Figure 5a bottom left; see Table S17 for details and other brain regions involved). Within each individual sub-group, we also found significant SPE signals in bilateral dlPFC within the MB group (cluster-level FWE-corrected, left dlPFC: *p* = 0.001; right dlPFC: *p* < 0.001), within the Mixture group (cluster-level FWE-corrected, left dlPFC: *p* < 0.001; right dlPFC: *p* = 0.012), but not within the MF group or the Other group (see Tables S18-S21 for all brain regions showing significant SPE signals in each sub-group). We then used an independent ROI of bilateral dlPFC based upon a previous finding of SPE encoding (Gläscher et al., 2010). Within this ROI, we found significant SPE encoding within each of the sub-groups, including the MF group (one-sample t-test: MB group, *p* < 0.001; Mixture group, *p* < 0.001; MF group, *p* = 0.0118; Other group, *p* < 0.001; Figure 5b). In a voxel-wise analysis, we also found significant SPE encoding around the area of IPS within the MB group (cluster-level FWE-corrected, bilateral IPS: *p* < 0.001), within the Mixture group (cluster-level FWE-corrected, left IPS: *p* < 0.001; right IPS: *p* < 0.001), within the MF group (cluster-level FWE-corrected, left IPS/superior parietal lobule/angular gyrus: *p* = 0.036, right IPS/superior parietal lobule: *p* = 0.001), but not within the Other group (see Tables S18-S21 for all brain regions showing significant SPE signals in each sub-group). When we used an independent ROI of bilateral IPS previously found to encode SPE (Gläscher et al., 2010), we found significant SPE encoding in the ROIs within each of the sub-groups, including the Other group (one-sample t-test: MB group, *p* < 0.001; Mixture group, *p* < 0.001; MF group, *p* = 0.0098; Other group, *p* = 0.0397; Figure 5d).

**Figure 5:**
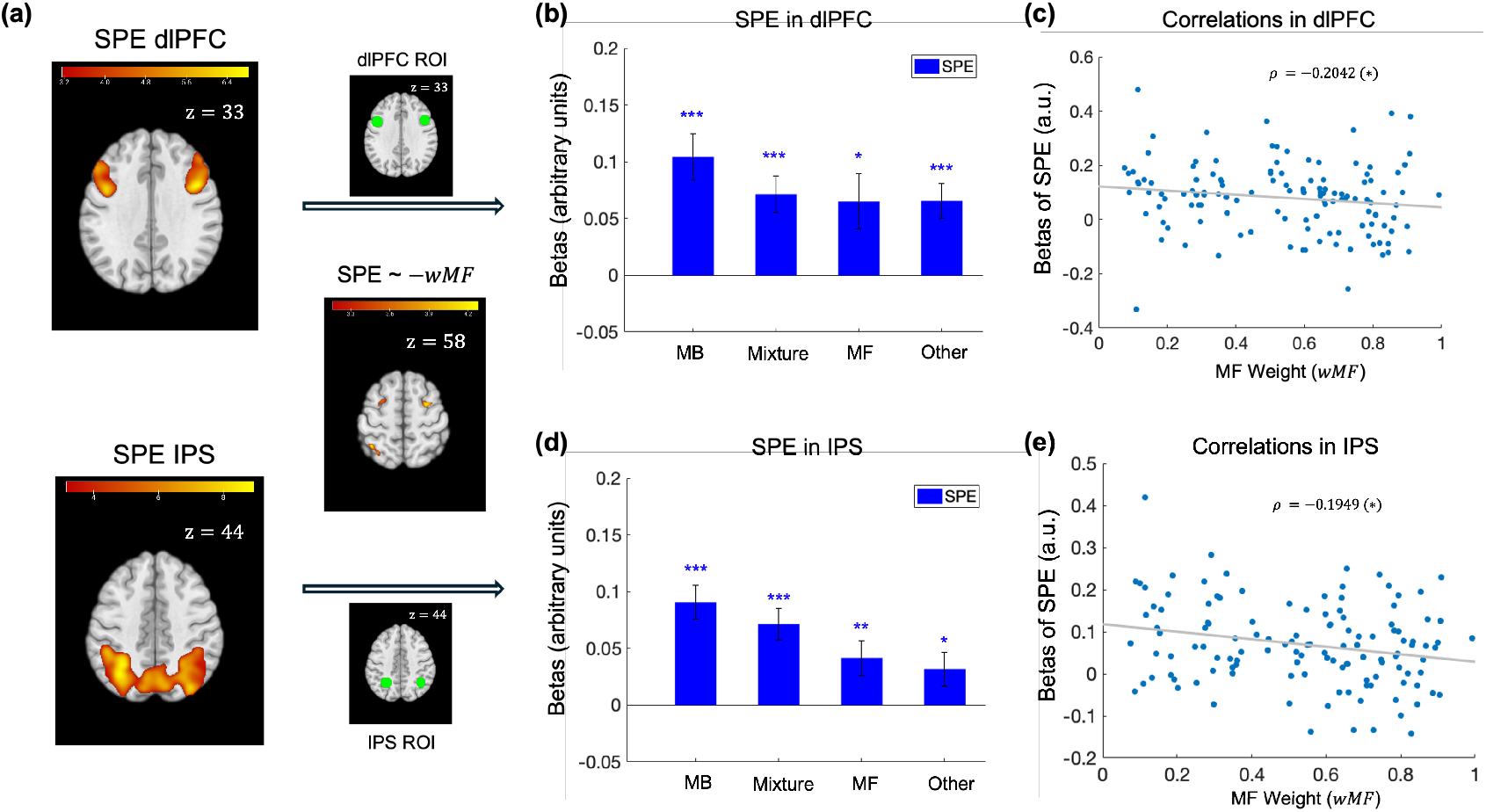
Encoding of state prediction errors at the group level, at the sub-group level, and across individuals. All brain images show the T-maps with a cluster-forming threshold of *p* < 0.001, and cluster-level FWE-corrected *p* < 0.05, in the corresponding regions of interest. (a) Top left: The clusters in the bilateral dorsolateral prefrontal cortex with activities correlating with SPE; Bottom left: The clusters in the bilateral intraparietal sulcus with activities correlating with SPE; Top right: The dlPFC mask used for ROI analyses of SPE betas in (b) and (c); Middle right: The clusters in the left intraparietal sulcus (IPS) and the bilateral superior frontal gyrus (SFG) where the encoding of SPE signals shows a significant correlation with the MB degree (i.e., −*wMF*); Bottom right: The IPS mask used for ROI analyses of SPE betas in (d) and (e). (b): The subgroup-level beta coefficients of the SPE contrasts in the ROIs of bilateral dlPFC (one-sample t-test). (c): The correlation between the beta coefficients of SPE in the ROI of the bilateral dlPFC and the weight of the MF system from the computational model (Spearman correlation, *: *p* < 0.05). (d): The subgroup-level beta coefficients of the SPE contrasts in the ROIs of bilateral IPS (one-sample t-test). (e): The correlation between the beta coefficients of SPE in the ROI of the bilateral IPS and the weight of the MF system from the computational model (Spearman correlation, *: *p* < 0.05). To denote statistical significance: ***: *p* < 0.001, **: *p* < 0.01, *: *p* < 0.05, n.s. : *p >* 0.05.

We also ran a second-level GLM analysis on how the SPE encoding degree was modulated by the derived *wMF* variable from the computational model, which indicates the behavioral MF tendency of a given participant. We found that the SPE encoding degree was negatively correlated with the *wMF* in left intraparietal sulcus (*p* = 0.013, cluster-level FWE-corrected, Figure 5a middle right), bilateral superior frontal gyrus (left superior frontal gyrus, *p* = 0.025, cluster-level FWE-corrected; right superior frontal gyrus, *p* = 0.014, cluster-level FWE-corrected; Figure 5a middle right; see Table S22 for details). In other words, the higher behavioral MB tendency a participant had, the stronger SPE encoding was in the aforementioned brain regions. To further investigate the relationship between the SPE encoding degree in our regions of interest and the degree of engaging MB control, we ran a regression model on the 1st-level beta estimates of SPE contrasts in both the dlPFC and IPS with an intercept and an ordinal group variable across three groups using the RL strategy (i.e., MB, Mixture, and MF groups). For both regions of dlPFC and IPS, we observed a significant overall SPE encoding, indicated by significant intercepts (dlPFC: *β* = 0.120, *p* < 0.001; IPS: *β* = 0.116, *p* < 0.001), and we also observed a significant categorical differences between the magnitude of the SPE signals in IPS across three RL subgroups and a trending effect was also observed in dlPFC (IPS: *β* = −0.0242, *p* = 0.0280; dlPFC: *β* = −0.0202, *p* = 0.1681). Yet, when we examined correlations between the SPE 1st-level beta estimates and the continuous MF weight (*wMF*) across individuals, we found significant correlations in both dlPFC and IPS ROIs (dlPFC: Spearman’s *ρ* = −0.2042, *p* = 0.0220, Figure 5c; IPS: Spearman’s *ρ* = −0.1949, *p* = 0.0289, Figure 5e), suggesting that across individuals using the RL strategies, the SPE encoding becomes stronger when MB control is more strongly engaged in behavior. These findings support the existence of SPE signals potentially underpinning learning of a state-action-state transition model even in participants who do not actively utilize an MB strategy, but also emphasize that the strength of SPE encoding varies as a function of the degree of expressed MB-consistent behavior.

We also would like to note that, to take into account the potential confounding effect of age on our findings, in an additional analysis we reran the 2nd level fMRI analyses while including age as a covariate for all relevant analyses on the state prediction error signal. The results presented in this section still hold up at the relevant statistical thresholds.

For both MB and MF RPE signals, we examined ventral and dorsal striatal ROIs, regions previously found to encode reward prediction errors (O’Doherty et al., 2003; Jessup and O’Doherty, 2011; Valentin and O’Doherty, 2009; McClure et al., 2003). We used a combined striatal ROI encompassing both the ventral and dorsal part of the striatum, which bilaterally covers nucleus accumbens, putamen and caudate (Pauli et al., 2018). When pooling across all participants with RL-consistent behaviors, although we found that a cluster in the right middle caudate was correlated with the combined MF RPE signal at the level *p* < 0.001 (uncorrected), the cluster did not survive cluster-level FWE-correction with small-volume correction using the combined striatal ROI (*p* = 0.195, cluster-level FWE-corrected, small-volume corrected, Figure S2 left). We also found that the neural activities of two clusters in the left and right ventral striatum respectively were correlated with the combined MB RPE signal at the level *p* < 0.001 (uncorrected), yet the two clusters did not survive cluster-level FWE-correction with small-volume correction using the combined striatal ROI (left ventral striatum: *p* = 0.130, cluster-level FWE-corrected, small-volume corrected, Figure S2 right; right ventral striatum: *p* = 0.246, cluster-level FWE-corrected, small-volume corrected). For MF and MB RPE, we did not find significant effects in these ROIs when testing for effects in each strategy sub-group separately, which likely reflects a lack of statistical power because of high intrinsic correlations between these two signals and between the RPE signals and the outcome magnitude regressor (mean Pearson correlations, MF RPE and MB RPE: 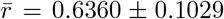;MF RPE and Outcome Magnitude 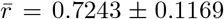; MB RPE and Outcome Magnitude: 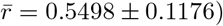.

## Discussion

We leveraged a large cohort of participants who underwent fMRI scans while performing a behavioral task designed to dissociate neural correlates of MB and MF computational variables in order to investigate the relationship between variation in model-based and model-free control expressed at the behavioral level across individuals, and underlying neuronal correlates of these strategies.

Consistent with prior reports (Hampton et al., 2006; Beierholm et al., 2011; Lee et al., 2014), we found robust evidence for MF and MB decision value signals in the vmPFC and a number of other brain areas when pooling over all participants, indicating that a mixture of MF and MB systems are engaged at the neural level across the whole sample, consistent with the mixture of strategies found to be expressed in behavior overall. However, there is considerable variability at the individual level insofar as which strategy is being utilized. While one subgroup of individuals indeed utilizes a mixture of MB and MF strategies, others were better characterized as either being predominantly MB, predominantly MF, or even found to be not well described by either strategy.

When focusing on distinct subgroups of individuals that were identified based on the predominant behavioral strategies they utilized, we found dissociable effects of MB and MF value signals across subgroups. When examining neural correlates of MB value across sub-groups within the vmPFC, we found that the MB value signal appeared to be directly sensitive to the degree to which MB control was present at the behavioral level. When plotted across the subgroups, and when examining correlations between the degree of MB control and the degree of MB value in vmPFC in each individual, we found a significant relationship between the amount of MB control exhibited behaviorally and the strength of the MB value representation in vmPFC and other brain structures. This finding supports the possibility that MB value signals are encoded in vmPFC and elsewhere only when they are being actively used to drive behavior.

The absence of a clear MB value signal in those who do not exhibit MB control at the behavioral level invites several interpretations. Firstly, these findings could simply reflect the fact that individuals (in the MF or other groups) did not actively perform MB computations at all but instead relied on a default MF or non-RL strategy, respectively. Yet another possibility is that individuals in the MF and other groups did indeed attempt to compute MB value signals but utilized either the wrong task model or were impaired at learning the correct task model. This latter explanation would also lead to MB value signals that are not detectable via our model-based regression analysis, which is conditioned on accurate task representations. As we discuss further down in more detail, our findings of reduced state prediction error signals in those with reduced MB control would appear to better support an account whereby individuals exhibiting reduced MB control are impaired at encoding and/or updating their world model, leading to reduced MB value signals being detected.

Most intriguingly, we did not observe a similar relationship between MF strategy use at the behavioral level and MF value signals as we did in the MB case. Unlike the case with MB value representations, MF value signals were ubiquitous in the brain across all sub-groups, irrespective of which strategy they used to control behavior. Most notably, both the MB and Other groups exhibited robust MF value signals in the vmPFC in spite of the fact that they exhibited little evidence of using this strategy in their behavior on the task (the Other group shows a somewhat blunted MF-like behavior). Thus, there is an asymmetry between MF and MB strategies, suggesting they are qualitatively different in how they are represented in the brain. The persistent presence of MF value signals across individuals could reflect the notion that MF value signals are a baseline default computation that occurs independently of behavioral control. Whether individuals utilize this MF strategy or not may depend on other factors such as arbitration trade-offs between MB and MF control and/or other strategies (Lee et al., 2014; O’Doherty et al., 2021). This could reflect the fact that MF value signals are relatively computationally straightforward to compute. More speculatively, perhaps MF RL is always present in the brain because it is an evolutionarily preserved reward-learning strategy that does not require a world-model or online planning to compute. One interesting observation was that the Other group showed a somewhat greater spatial extent of activation for MF decision value signal compared to the MF group. We would like to note that first of all, observing significant MF decision value signal in the Other group is reasonable as a majority of the Other group’s participants (31 out of 53) is best explained by the MF model than any other models (such as arbitration/fixed-weight mixture models, MB model or random model, *p* < 0.001, two-proportion Z-test). The Other group showed a better-than-chance performance in collecting rewards and a lower softmax inverse temperature than the MF group (with comparable learning rates), indicating a somewhat blunted MF strategy in use, which would suggest the MF decision value computations were still carried out by the Other group. Thus, the larger group size of the Other group (N=53) compared to the MF group (N=34) would possibly contribute to a more robust estimation of MF decision value thereby eliciting a more robust activation profile that we observed here.

Beyond the value signals carried by these two systems, we also tested for neural correlates of prediction errors, the putative signals involved in mediating learning of relevant representations. One important signal in model-based learning is the state-prediction error. This signal is proposed to mediate learning of state-action-state transitions that underpin the cognitive model utilized by the model-based system (Gläscher et al., 2010). Consistent with prior studies, we found evidence for state prediction error signals in the posterior parietal cortex and dorsolateral prefrontal cortex (Gläscher et al., 2010; Lee et al., 2014), supporting the role of dorsal cortex in implementing learning of stateaction-state transitions relevant for model-based control. The magnitude of these error signals was found to be related to the degree to which behavior is model-based. SPE signals were stronger in those with greater MB control behaviorally, and weaker in those with more MF control, in both parietal and prefrontal cortex regions. This finding supports one of the possible explanations outlined earlier for reduced model-based value control and reduced model-based value signals. Reduced SPE signals in those with reduced model-based control could indicate that participants have failed to effectively learn an accurate model of state-action-state transitions, because their SPE signals are either weaker overall or being computed incorrectly. Thus, reduced model-based control may have occurred because of a reduced capacity to either encode or learn about a model of the world. If individuals lack robust knowledge of the transition structure, then they may struggle to compute accurate model-based value signals. Consequently, the finding of reduced model-based value representations in vmPFC may relate to alterations in the capacity to compute and update the cognitive model used to underpin model-based inference.

Lastly, one aspect of the behaviors exhibited in the current sample is that participants used a variety of behavioral strategies, ranging from MF, MB, a mixture of MF and MB, and non-RL-like behaviors. One possible contributing factor to the heterogeneity in strategies observed is heterogeneity in the population drawn from: the present study involved participants recruited from a general population of Los Angeles community members as opposed to individuals drawn from a predominantly university campus setting. This may have contributed to a greater variety in the strategies seen than would be the case had we relied on a predominantly university-based sample. On the other hand, there has been some discussion on the effects of task complexity in the two-step task and how this might have a potential confounding effect on the use of behavioral strategies in the task and on the identification of the true model underlying the expressed behaviors (Akam et al., 2015). We show through simulations that even though the present task might be considered to possess complex elements, it is not vulnerable to a problem such as model misidentification, and furthermore, the expressed RL behaviors are well explained by their corresponding RL models (Figure S4).

To conclude, the present results provide insight into the relationship between model-based and model-free computations in the brain and their relationship to individual differences in model-based and model-free behavior. While MF value signals appeared to be present in the vmPFC irrespective of whether or not those signals are being actively utilized to drive behavior, MB signals in the vmPFC are directly sensitive to the degree to which those signals are being used to actively drive behavior within an individual. Furthermore, we find evidence for distinct MB and MF reward prediction error signals within the ventral and dorsal striatum, respectively. When taken together, these findings reaffirm the existence of parallel and distinct model-based and model-free learning systems in the brain, while providing empirical support for the ubiquity of model-free learning, even in those who do not utilize a model-free strategy at the behavioral level.

## STAR Methods

### Resource Availability

#### Lead Contact

Requests for further information and resources should be directed to and will be fulfilled by the lead contact, Dr. John P. O’Doherty.

#### Materials Availability

This study did not generate new unique reagents.

#### Data and code availability

Neural data reported in this paper will be available from this link on OpenNeuro:

https://openneuro.org/datasets/ds007474.

Behavioral data will be available at this link on OSF:

https://osf.io/ctfzd/overview?view_only=022584fa16944755982f0d06a713835c.

Code will be available at this GitHub repository:

https://github.com/Gentu-Ding/2-step-task-MBMF-fMRI-code.git.

Any additional information can be requested via contact.

### Participants

#### Recruitment and Inclusion

We recruited participants from the greater Los Angeles area who are fluent English speakers and readers. The age range for the studies is from 18 to 65 years. Before the experiment, all participants signed the informed consent approved by the California Institute of Technology’s Institutional Review Board under protocol 19-0914. All participants were reimbursed in monetary form for their base pay and their performance bonus. There were originally a total of 189 participants who completed the two-step task with functional magnetic resonance imaging data. After data exclusion based on head motion (see the next section on the Exclusion Criteria), 179 participants (105 females) were retained, with a mean age of 30.37 years (*sd* = 10.19). We conducted pre-screening of the participants so that participants in the experiment did not report any history of substance/alcohol use disorder, anxiety disorder (Obsessive-Compulsive Disorder, Body Dysmorphic Disorder, generalized anxiety, social anxiety/social phobia), and/or depressive disorders (dysthymia, major depression). Furthermore, participants reported they did not use any medications for subclinical psychiatric disorders.

#### Exclusion Criteria

Among the original 189 participants, we excluded the participants’ data based on the proportion of fMRI volumes that exceeded a manually specified motion threshold to eliminate potential neural confounds due to too much head motion. Ten participants were excluded from this procedure, giving us 179 participants’ data in the main analysis. Participants who had at least one fMRI run with 15% of the total volumes exceeding the framewise displacement (FD) threshold of 0.77mm were excluded. The threshold of 0.77mm is determined by first calculating the FD threshold of 1.5 times the interquartile range plus the third quartile of each participant’s FD across all scans, then performing the calculation of 1.5 times the interquartile range plus the third quartile of the entire FD threshold distribution across all individuals.

We also want to note that there are five participants who only experienced the state-contingent reward condition but not the stimulus-contingent reward condition due to an experimental operation error. We still included these five participants in our main analysis, as our focus here is not the effect of the contingency manipulation but the overall value and error signals throughout the task.

### Experimental Procedure

Participants first gave informed consent. The participants were then screened through a structured clinical interview conducted by a psychiatrist, in which their mental health and medication usage were evaluated to determine their eligibility to participate as healthy controls, psychiatric patients, or to be excluded from further participation. In the present study, we report only on the data from healthy participants. The data from psychiatric patients will be reported elsewhere and is beyond the scope of the current study.

Participants were also asked to complete a few questionnaires on psychiatric measures before coming to the Brain Imaging Center at the California Institute of Technology for MRI scanning.

Firstly, upon arrival at the Brain Imaging Center, participants were asked to complete a short questionnaire asking about feelings and emotions. Afterwards, before the fMRI scan, participants read through the instructions for the spaceship task (a variant of the original two-step task). After the instructions, several questions were asked to ensure participants understood the task and remembered the key features of the task. After the instructions, 115 trials of practice rounds of the task were done before participants performed the main task in the scanner. The spaceship task was paired with another task, studying habits, and the participants completed both tasks during the scan. The MRI scans were split into two half-sessions where participants completed one task per session; completion of the spaceship task in the first or the second half was counterbalanced across participants. Data from the habit task will also be reported elsewhere and is beyond the scope of the present study.

During the structural scan in the scanner (either in the first or the second half session), participants also completed another session (105 trials) of the spaceship task right before the functional session (154 trials). Task behaviors during the structural scan and the functional scan were combined for the analysis of behavior on the task, as well as for the computational modeling of the behavior, so that a maximal number of trials could be used. After the scanning procedure was completed, participants were debriefed about the task and received their reimbursement.

### Space Miner Task

Participants were instructed to collect rewards through space mining on different planets. The back-story for the task is that mining has begun on two planets (i.e., one identified as red and the other green) in space, and the participants’ goal is to earn as many points as possible by mining gems from the two planets. However, the mines on the two planets have changing conditions in their production. Sometimes gems could be found, but other times, worthless rocks could also be mined out. Specifically, there are two landing pads for the corresponding mines on each planet, one to the North and the other to the South. The landing pads are identified through their unique scenic view and are located in the upper (North) and bottom (South) parts of the planet on the screen.

Participants can choose between two different spaceships (identified as yellow and blue, with the screen locations of the spaceships fixed across trials) to travel to the two planets for mining. They are instructed that the yellow spaceship usually lands on the red planet, and the blue spaceship usually lands on the green planet. However, space travel can sometimes be a bit unpredictable due to space debris, so in some rare situations, the yellow ship will be forced to land on the green planet, and the blue spaceship will be forced to land on the red planet. Participants were instructed to press one of two buttons to choose the corresponding spaceship. In a given trial, once participants chose one spaceship, they observed the spaceship being highlighted and taken off, and the planet appearing after the spaceship landed. Afterwards, the landing pad for the mine would also appear, either on the upper or the bottom part of the planet, depending on whether they landed at the North or the South mine on the planet. Once the spaceship landed, the mine production appeared as either a gem with its price or a worthless rock. Specifically, participants were instructed that the gem’s price is unpredictable and will change daily. Participants had no control over which mine (North or South) the spaceship would eventually land on. In general, the conditions at all four mines (two mines on each of the two planets) changed throughout the game regarding gem vs. stone production. Participants were encouraged to learn which mine produces gems the most reliably.

After going through the instructions of the spaceship task, participants were asked a few questions:

1. How many planets are there?
2. How many mines are there on each planet?
3. Which planet does the yellow ship usually land on?
4. Which planet does the blue ship usually land on?
5. How many points is a mined rock worth? (0 points vs. 1-100 points)
6. How many points is a gem worth? (0 points vs. 1-100 points)

Participants proceeded to the practice trials and then the main experiment if they answered all the questions correctly.

The task was programmed using Psychtoolbox-3 in Matlab (Brainard and Vision, 1997; Pelli, 1997; Kleiner et al., 2007).

### Task Design

The current variant of the two-step task (Cockburn et al., 2024) shared a similar task structure as the original two-step task (Daw et al., 2011) overall, but with some differences in details. The general structure of the present task consists of the first-step transition and reward probability shifts, alongside three condition manipulations (i.e., reward magnitude, state transition, and reward contingency). One key difference between the present task and the traditional two-step task is that in the current task, there was only one action to be made per trial, which was at the initial state, while the rest of the trial was all state transitions in which no further actions were required. On each trial, the yellow and blue spaceships would appear on the left and right sides of the screen. If chosen, the transition via the yellow spaceship towards the red planet occurred with a probability of 0.7 and with a probability of towards the green planet; the same probabilities apply for the transitions from the blue spaceship to the green and red planet, respectively. After the transition from the planet stage to the landing-pad stage, reward or non-reward outcomes would appear. The underlying reward probability was shared across the two landing pads on a given planet (reward probability was also associated with the spaceship chosen, depending on the reward contingency condition described later in this section). There were specific periods built in such that landing on one planet was more rewarding than landing on the other planet, and also periods where landing on either of the two planets was relatively comparable in terms of the reward probability. To implement this, the reward probability associated with the two planets started from 1 vs. 0, and then by using a Sigmoid function, the reward probability of the rewarding planet decayed from 1 towards 0.3 (asymptote) within the time span of from 20 to 25 trials (the exact number of trials depended on whether a rare spaceship-planet transition is made); for the reward probability associated with the current non-rewarding planet, the same Sigmoid decay rate with a flipped sign for the decay slope was used to set the reward probability drift from 0 to 0.5 (asymptote) again within the time span of from 20 to 25 trials. After this, the reward probabilities associated with the two planets were reset to 1 vs. 0, but the rewarding planet was reversed compared to the previous block of trials. This drifting of reward probabilities and the reversal of rewarding planet/landing pads occurred throughout the entire session of the experiment. The order of which planet was first rewarding was counterbalanced across the participants. The structure of going from a strong preference (large gap of reward probabilities: 1 vs. 0) towards almost indifference (reward probabilities: 0.3 vs. 0.5) was first to facilitate learning of the rewarding option and then for setting the value of two options towards indifference to prepare for learning after the reward probability reversal. Also, a rare transition occurred immediately after a reward probability reversal to facilitate detecting stay/switch behaviors as a signature of the MB vs. MF strategy. It is worth noting that another manipulation of reward contingency (described later in this section) would change the contingency of reward delivery upon the landing pads (or planets) vs. the spaceships, yet the general fluctuating and reversal dynamics of the reward probability would remain the same across the two reward contingency conditions.

On top of the reward probability shifts that occur throughout the main task, three additional manipulations were included: 1. Manipulations of reward magnitude (low and high switching every 26-27 trials) with the reward distribution varying from 0.1 to 0.19 for low, and from 0.3 to 1 for high, for which the actual points were scaled by 100. 2. Manipulations of state-transition uncertainty from the planet to the landing pads where during the high state-transition uncertainty condition, the transition to one of the two landing pads on a given planet was 0.5 vs. 0.5, while during the low state-transition uncertainty condition, the transition to one of the two landing pads on a given planet was 0.9 vs. 0.1 (also switching every 26-27 trials offset w.r.t. to the reward magnitude manipulation). 3. Manipulations of reward contingency where the reward delivery was contingent upon either what stimulus (i.e., spaceship) participants choose or what terminal states (i.e., landing pads) were arrived at, meaning rewards are either stimulus-contingent or state-contingent, respectively. The stimulus-contingent and state-contingent conditions were run in two separate fMRI blocks of the experiment, and the two fMRI blocks had the same number of trials (77 trials each, 154 trials in total), and the order of the two contingency conditions was also counterbalanced across participants. More details about these manipulations and their rationale are provided elsewhere (Cockburn et al., 2024). The present study does not focus on the effects of these conditional manipulations, given that the present study is focused on the topic of between-subject variation in overall model-based and model-free behavioral strategies, while these manipulations are focused on within-subject shifts in behavioral control. The effects of these manipulations will be described in a separate manuscript. However, trial-by-trial variation in the task variables introduced via these manipulations is implicitly captured in the computational model described below, because the computational model fit to behavior and used to generate regressors for the model-driven fMRI analysis takes into account shifts in control within each subject across conditions, so we could more precisely capture the task behavior within each participant.

### Statistical Analysis of Behavior

#### Logistic Regression

For all the mixed-effect logistic regression analyses, the “fitglme” function in Matlab was used. To quantify how the previous reward, previous transition type, and their interactions affected the stay choices, we used a mixed-effect logistic regression to model the probability of staying with the option chosen in the previous trial *isStay*_*t*_ (1: stay, 0: switch) as a function of the previous outcome *Reward*_*t*−1_ (1: reward, -1: no-reward), the previous transition type *Rare*_*t*−1_ (1: rare, -1: common) and their interactions. The fixed effects include the previous outcome, the previous transition type, and their interactions. The intercept and the slope of the previous outcome, the previous transition type, and their interaction were modeled as random effects that could vary at the single-subject level (indicated by *subID*). The regression model is specified as:

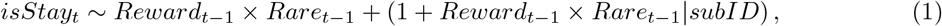

where “ ×” denotes the main effects and the interaction between each independent variable, and “1” denotes the intercept, which is the average stay probability for each subject. Trials, where the choice was not made within 2 seconds, were excluded before estimating the regression model.

#### Linear Regression

For the statistical test on the linear relationship between the random effects of outcome/outcome-transition and the behavioral MB (or MF) tendency entailed by the group identity obtained from the clustering, a linear regression model was run on the outcome random effects with an intercept and an ordinal group variable:

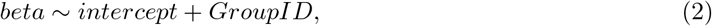

where, when testing for the outcome effect, “*beta*” denotes the random effects of the outcome for all participants excluding the Other group, and the ordinal variable “*GroupID*” has the coding of “1” for the MB group, “2” for the Mixture group, and “3” for the MF group; when testing for the outcome-transition effect, “*beta*” denotes the random effects of the outcome-transition for all participants excluding the Other group, and the ordinal variable “*GroupID*” has the coding of “1” for the MF group, “2” for the Mixture group, and “3” for the MB group.

### Behavioral Clustering

There are, in general, two types of behavioral metrics we leveraged to perform cluster allocation on participants’ behavior: 1) choice measure: choice pattern as a function of the outcome and transition type in the preceding trial, and 2) reaction time measure: reaction time pattern as a function of current choice, the outcome and transition type in the preceding trial. Specifically, within the choice measure, summary statistics and regression estimates of each participant served as the features for the clustering. For the summary statistics, we calculate the following metrics for each participant:

1. *p*(*Stay*_*t*_|*Common*_*t*−1_*Rewarded*_*t*−1_)
2. *p*(*Stay*_*t*_|*Common*_*t*−1_*Unrewarded*_*t*−1_)
3. *p*(*Stay*_*t*_|*Rare*_*t*−1_*Rewarded*_*t*−1_)
4. *p*(*Stay*_*t*_|*Rare*_*t*−1_*Unrewarded*_*t*−1_)
5. Reward Sensitivity:

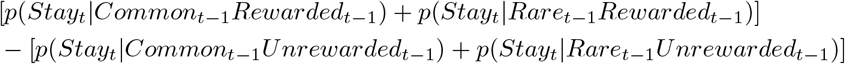
6. Transition Sensitivity:

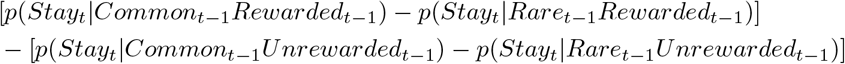
7. Absolute Reward Sensitivity:

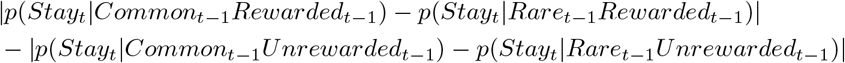

For the regression estimates of the choice measure, we used a mixed-effect logistic regression to model the choice repeating probability as a function of the intercept, outcome, transition type, and their interaction in the preceding trial, which serve as the fixed effects. The dependent variable “*isStay*_*t*_” denotes whether choosing the same option as in the previous trial, and all regressors are dummy variables identifying the status in the preceding trial: “*Reward*_*t*−1_” denotes the outcome in the previous trial; “*Rare*_*t*−1_” denotes the spaceship-planet transition type in the previous trial. Three binary variables indicating the conditional manipulations are added as three independent interactions with “*Reward*_*t*−1_ × *Rare*_*t*−1_”. Specifically, “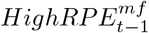 “ is a binary variable indicating the previous trial’s reward magnitude condition (1: high reward magnitude/high MF-RPE, -1: low reward magnitude/low MF-RPE). Similarly, “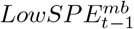 “ indicates the previous trial’s state-transition uncertainty condition (1: low state-transition uncertainty/low SPE, -1: high state-transition uncertainty/high SPE), and “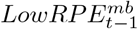” indicates the previous trial’s reward contingency condition (1: state-contingent reward/low MB-RPE, -1: stimulus-contingent reward/ high MB-RPE). For the random effects, the intercept, the slope of the outcome, transition type, their interaction in the preceding trial, and the independent interactions of three conditional manipulations with “*Reward*_*t*−1_ ×*Rare*_*t*−1_” are also modeled to vary at the single-subject level. And “subID” denotes the identity of each participant. Hence, the full model is:

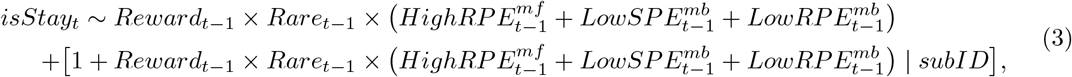

where “ ×” denotes the main effects and the interaction between each independent variable, and “1” denotes the intercept, that is, the average stay probability for each subject. Trials, where the choice was not made within 2 seconds, were excluded before estimating the regression model. From the regression, we extracted the random effects of the intercept, the outcome, the transition type, and the outcome-transition interaction from each individual to use as features in the clustering algorithm.

Regarding the reaction time measure, we also used a mixed-effect logistic regression to characterize how individuals’ reaction times (RT) varied as a function of the current choice *Stay*_*t*_ (1: stay, -1: switch), the preceding trial’s outcome *Reward*_*t*−1_ (1: reward, -1: no-reward), the preceding trial’s transition *Rare*_*t*−1_ (1: rare, -1: common), and all the two-way and three-way interactions between these regressors. The same were modeled as random effects for each individual.

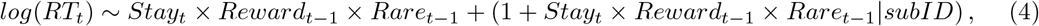

where we log-transformed the RT, and “ ×” denotes the main effects and the interaction between each independent variable, and “1” denotes the intercept. Trials, where the choice was not made within 2 seconds, were excluded before estimating the regression model. We also extracted each individual’s random effects of the intercept, the outcome, the transition type, and the outcome-transition interaction for their clustering features.

In total, from both the choice measures (summary statistics and regression estimates) and the reaction time measures (regression estimates), there are 19 behavioral features for each participant. We relied on the centroids discovered in an external dataset with a much larger sample size (Cockburn et al., 2024) to perform cluster allocation for this dataset. In the external dataset, a much larger group of participants (N=678) completed the same two-step task online, and a k-means clustering algorithm was used on the same 19 behavioral features to classify all the participants, and clusters’ centroids were obtained. The optimal number of clusters was found to be 4 using the Gap statistical method (Tibshirani et al., 2001) in the external dataset, which gave sensible classification to separate behaviors generally as RL strategies vs. non-RL strategies as well as providing finer-grained strategy characterization for participants using RL strategies - 1) A Mixture of MF and MB control, 2) pure MF control, and 3) pure MB control. In the current dataset, we assign each individual to the group label of the closest centroid in terms of Euclidean distance. The labels were defined according to behavioral phenotypes in the external dataset: 1) Mixture Group, 2) MF Group, 3) MB Group, and 4) Other Group (non-RL strategies).

### Computational Models

We used an arbitration mixture model (Arbitration Mixture), a hybrid reinforcement learning model composed of a model-free module and a model-based module, to characterize the participants’ behaviors in this task and extract computational variables for the fMRI analysis. We will describe the MF module and the MB module of the mixture model, and how they are integrated to make the arbitration mixture model (Arbitration Mixture). We will also briefly describe other candidate models considered in the model comparisons: 1) the model-free model (Model-Free), 2) the model-based model (Model-Based), and 3) the fixed-weight mixture model (Fixed-weight Mixture), all of which are nested models of the Arbitration Mixture model.

#### Model-Free Module

The model-free module is composed of a slow learning and a fast learning component, which corresponds to the learning over outcomes of multiple past trials (“slow”) and learning upon the outcome of the preceding trial (“fast”). The slow learning component is described first.

As the space miner task only requires one action to be performed at the initial stage, leading to two subsequent intermediate states before the outcome is revealed, the MF module only learns the action values of the two stimuli at the initial stage. As the position of the two stimuli at the initial stage is fixed, choosing the left (or right) action always led to the yellow (or blue) spaceship, and thus the action values learned were effectively the same as the stimulus values. In other words, the MF module essentially learns the value of potential states 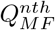 (*s*) (*n* as the stage order in a sequence, *s* as the potential states within the nth stage). Hence, for example, the Q-value of the yellow spaceship could be denoted as 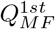 (*yellow*), and the Q-value of the red planet and the Q-value of the north pad on the red planet could be denoted as 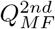 (*red*) and 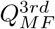 (*red* -*north*), respectively. The reward prediction error experienced at the nth stage could be denoted as 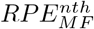 (*n* = 2 when at the planet stage, *n* = 3 when at the landing pad stage, and *n* = 4 when observing the outcome).

The reward prediction error at each stage and the learning rule for Q-values in previous stages (with eligible traces) were defined as the following (the *n* indicates the stages to which the corresponding equation applies; the subscripts of *MF* and the chosen/experienced state variable *s* for Q-values and *RPE* are not shown for simplicity):

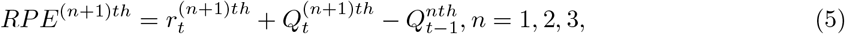

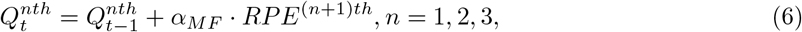

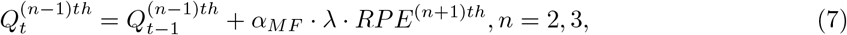

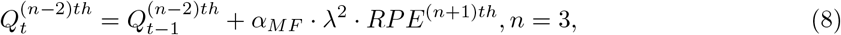

where the *α*_*MF*_ (0 *< α*_*MF*_ *<* 1) is the learning rate of the Q-values and shared across all stages, *n* = 1, 2, 3, within the trial; the trace decay parameter *λ* is set to be 1, meaning that an equal amount of value updates was conducted to a proximal and to a distal state. All-stage Q-values at the beginning of the session were initialized to 0 (*n* = 1, 2, 3). 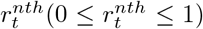 denotes the magnitude of the outcome (scaled down by 100) and is only relevant when *n* = 3, which is when observing the actual outcome, but otherwise 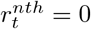 given no reward delivery in the intermediate states. Note that 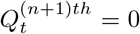 when *n* = 3, as there are no learnable states besides the outcome at that stage. Values of non-visited states or the unchosen stimulus are not updated and stay the same as in the previous trial.

As for the fast learning component, a value bias *W*_*t*_(*s*_*t*−1_, *r*_*t*−1_) was added to the learned Q-values of the preceding trial’s first-stage chosen stimuli *s*_*t*−1_, dependent on the outcome of the preceding trial *r*_*t*−1_ ∈ {0 : *no reward*, 1 : *reward}*. It essentially builds a stay or a switch action tendency bias into the associated stimulus based upon a preceding reward or no-reward event. The incorporation of the fast learning component into the overall value of the first-stage chosen stimulus 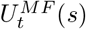 (*s*) is specified as follows:

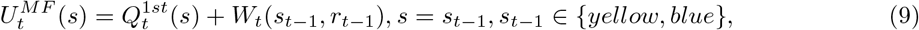

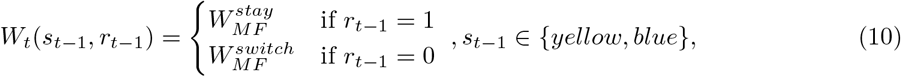

where 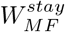 and 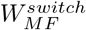 are the two parameters fitted for each participant, and both parameters could be positive or negative to capture all possible action adjustment policies. The value of the unchosen stimulus (*s* ≠ *s*_*t*−1_) would not be updated in the “fast” learning process and would stay the same as the stimulus value in the previous trial:

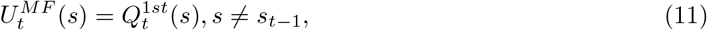

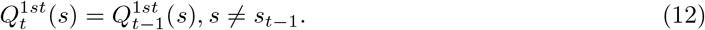

#### Model-Based Module

The model-based module is a forward learner using the dynamic programming approach, and it also consists of a slow learning component that updates the Q-values on a multi-trial basis and a fast learning component that considers the trial experience in the most recent trial. The slow learning component is first described below.

As described for the model-free module, since there is only one action needed in the task and the action values at the initial state are equivalent to the stimulus values, the Q-value learned in the MB module could be similarly denoted as 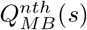 (*s*) (*n* as the stage order in a sequence, *s* as the potential states within the nth stage, as listed for the MF module). To carry out dynamic programming to compute the Q-values of upper-level states, the reward information (binary) was essentially learned in the MB module through 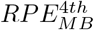 at the outcome stage for the Q-value of the experienced landing pads 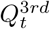:

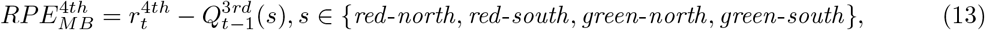

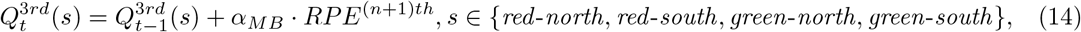

where *α*_*MB*_ (0¡*α*_*MB*_¡1) is the learning rate for the MB module. 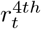 is a binary variable of 1 when rewarded and 0 when unrewarded in the current trial. Values of non-visited states are not updated and stay the same as in the previous trial.

Besides learning the binary reward information, the MB module also includes a magnitude learning component to incorporate the learned magnitude of the reward experienced at a certain landing pad *Mag*(*s*). The learned magnitude component *Mag*(*s*) is set to zero at the beginning and updated through a prediction error signal, only when a reward is encountered:

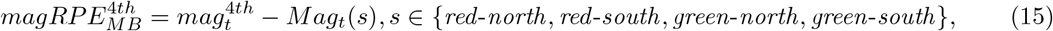

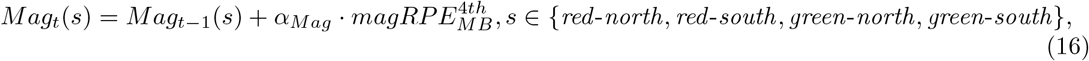

where 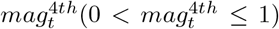 denotes the reward magnitude encountered, and *α*_*Mag*_ denotes the learning rate for the reward magnitude, and is set to 1 in the model.

When there is no reward, the magnitude component *Mag*(*s*) is set as:

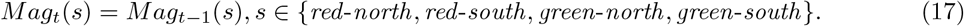

For non-visited terminal states, the learned magnitudes *Mag*_*t*_(*s*) for those states are not updated and stay the same as in the previous trial.

Then the value of the terminal states *Q*^3*rdMag*^(*s*) in the current trial is computed by adding the learned reward magnitude at the landing pads *Mag*(*s*) to the value component *Q*^3*rd*^(*s*) (learned through the binary reward information):

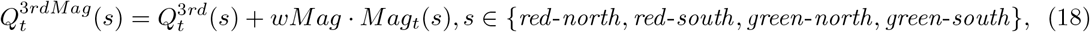

where *wMag*(*>* 0) is an unbounded positive parameter to set the weight of the learned magnitude component in value addition.

The MB module utilizes two sets of state-transition probabilities to compute the upper-level Q-values from the learned Q-value of the landing pads. The first state-transition probability is set as fixed instead of through learning, as assumed by the fact that participants were instructed about the common vs. rare transition structure from the spaceship and the planet, and had enough time to practice before the main experiment. The first state-transition probability *T*_1_(*s, s*^*′*^), *s* ∈ *{yellow, blue}*, and *s*^*′*^ = *{red, green}*, denotes the probability of transitioning from *s* to *s*^*′*^, and is specified as below:

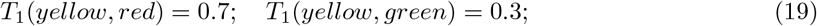

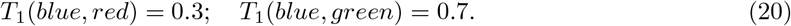

The state prediction error experienced during the first transition (spaceship to planet) is computed as:

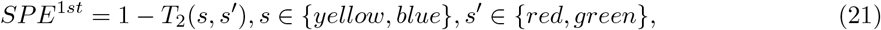

In contrast, the second set of state-transition probabilities *T*_2_(*s, s*^*′*^), ∈ {*s red, green}*, and *s*^*′*^ ∈ {*north, south}* is learned through state prediction errors (*SPE*) experienced through the transition from state *s* to the subsequent state *s*^*′*^ (i.e., from a planet to either the north or the south landing pad on that planet). The second state-transition probability *T*_2_(*s, s*^*′*^) is updated via the state prediction error experienced in the second transition (planet to landing pad) as follows:

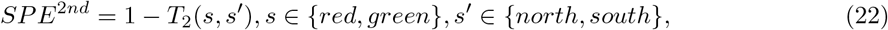

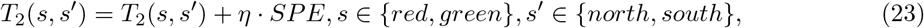

where *η* = 0.5 and it is the state-transition learning rate, and is a fixed value, as it is not a recoverable parameter through model fitting, and thus set as a median value of 0.5. For the other non-experienced state *s*^*′′*^, the transition probability was scaled by (1-*η*) to normalize the transition probabilities associated with the state *s*:

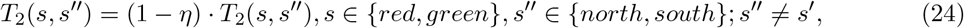

With the Q-value of the terminal states after the incorporation of the learned magnitude component 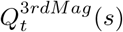, and the first and second sets of state-transition probabilities, the Q-values of the planets and that of spaceships 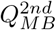 could be computed through dynamic programming as below (the subscripts *MB* are not shown for simplicity):

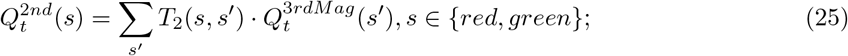

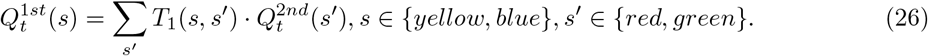

In addition to the slow learning component, the MB module also integrates information from the preceding trial to rapidly adjust the action tendency of staying with the selected option versus switching to the other option, which is referred to as the fast learning component. Specifically, to facilitate the stay vs. switch action selection tendency, the fast learning component utilizes the task model, considering the outcome and the first-stage transition type in the preceding trial, to add either a positive or negative value bias *W*_*t*_(*s*_*t*−1_, *r*_*t*−1_, *c*_*t*−1_) to the option selected in the preceding trial, where *s*_*t*−1_ ∈ {*yellow, blue}* denotes the selected spaceship, *r* ∈ {0 : *no*-*reward*, 1 : *reward}* denotes the outcome, and *c* ∈ {0 : *rare*, 1 : *common}* denotes the transition type. The value integration of slow and fast learning into the overall value of the first-stage chosen stimulus 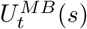 is specified as follows:

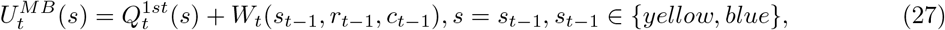

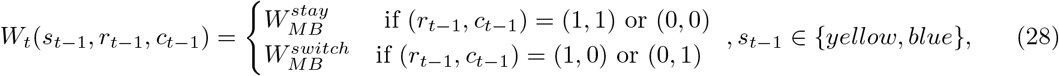

where 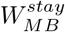 and 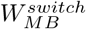 are the two parameters fitted for each participant, and both of the parameters are unbounded and could be either positive or negative to capture all possible action adjustment policies. The value of the unchosen stimulus (*s* ≠ *s*_*t*−1_) would not be updated in the “fast” learning process and would stay the same as the stimulus value in the previous trial:

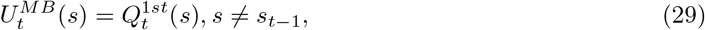

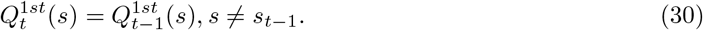

### The Arbitration Mixture Model with the MF and MB Modules

With the learned first-stage stimulus value from both the MF and MB modules, integration of the utilities from the two modules was implemented to obtain the mixed value of first-stage stimulus, which was used for action selection. The learned MF stimulus value 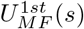 and MB stimulus value 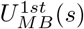 are combined through a weighting parameter *wMF* that decides the weight of value from the MF module for combination, and (1− *wMF*) is the weight of the MB stimulus value. A semi-arbitration mechanism is introduced to calculate the value of *wMF* for trials within certain reliability conditions. Considering the three manipulations of the prediction error signal throughout the experiment, as hypothesized by the reliability-based arbitration theory (Lee et al., 2014), we set three free parameters that correspond to the weight adjustment of the MF system in response to the three manipulations:

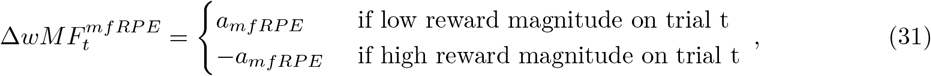

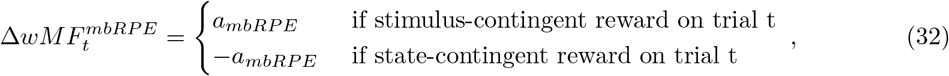

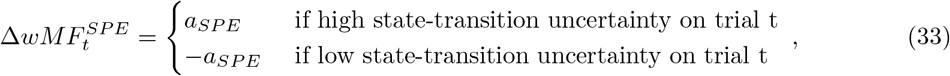

where *a*_*mfRP E*_, *a*_*mbRP E*_, and *a*_*SP E*_ are the three free parameters to fit. Based upon what the manipulated reliability condition a given trial *t* is in, the weight adjustments are integrated into the baseline weight of the MF module:

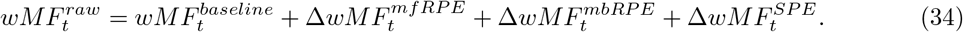

The adjusted raw MF weight is then passed into the sigmoid function to transform the raw weight onto the scale of (0,1):

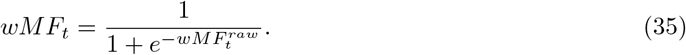

The weight assigned to the MB module on trial *t* would then be 1 −*wMF*_*t*_. Hence, the mixture of the MF stimulus value and the MB stimulus value is achieved by:

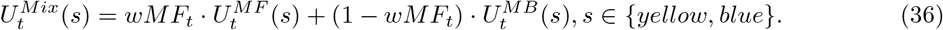

Once the mixed stimulus value is computed, the probability of choosing a specific stimulus *s* is then calculated through a softmax function:

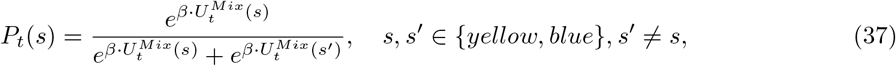

where *β* denotes the inverse temperature With the calculated choice probability for each participant, we fit the specified parameters to maximize the summed negative log-likelihood of participants’ choices across all trials (with no-response trials excluded):

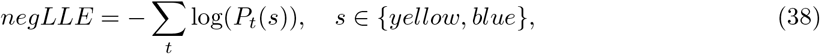

where *s* denotes the participant’s actual choice on trial *t*.

For each individual, the model parameters were fitted with a Bayesian inference method using the cbm (computational/behavioral modeling) toolbox (Piray et al., 2019) with the non-hierarchical specification. Each parameter was fit using a normally distributed prior with a mean of zero and a variance of 6.25 that ensured the coverage of a large range of parameters with no excessive model complexity penalty.

### The Model-Free Model, the Model-Based Model, and the Fixed-weight Mixture Model

The model-free (Model-Free) and the model-based (Model-Based) models correspond to the MF and MB modules, respectively, described in the previous subsection.

The fixed-weight mixture model (Fixed-weight Mixture) has the same structure as the arbitration mixture model (Arbitration Mixture) outlined in the previous subsection, except that the three weight-adjusting parameters take on the value of zero. Consequently, the model maintains a fixed degree of MF (or MB) tendency captured by the free parameter *wMF*^*baseline*^ throughout the task:

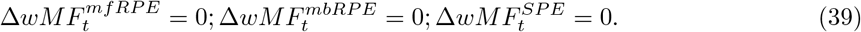

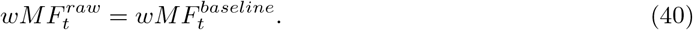

Hence, the Fixed-weight Mixture model has three fewer free parameters than the Arbitration Mixture model.

The Model-Free, Model-Based and Fixed-weight Mixture Models computes the probability of choosing a certain stimulus through the same softmax function on the learned 1st-stage stimulus values, as described for the Arbitration Mixture model:

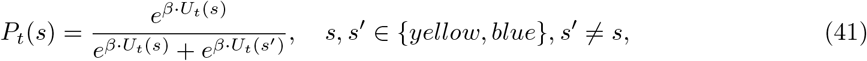

where *U*_*t*_(*s*) denotes the first-stage stimulus value for each of the corresponding model.

The same model fitting procedure for the Arbitration Mixture model was applied to these three candidate models.

### Functional MRI Acquisition

The fMRI data were acquired at the Caltech Brain Imaging Center (Pasadena, CA) using a Siemens Prisma 3T scanner with a 32-channel radiofrequency coil. The functional scans were conducted using a multi-band echo-planar imaging (EPI) sequence with 72 slices, -30 degrees slice tilt from AC-PC line, 192 mm × 192 mm field of view, 2 mm isotropic resolution, repetition time (TR) of 1.12 s, echo time (TE) of 30ms, multi-band acceleration of 4, 54-degree flip angle, in-plane acceleration factor 2, echo spacing of 0.56 ms, and EPI factor of 96. Following each run, both positive and negative polarity EPI-based field maps were collected using similar parameters to the functional sequence but with a single band, TR of 5.13 s, TE of 41.40 ms, and 90-degree flip angle. T1-weighted and T2-weighted structural images were also acquired for each participant with 0.9 mm isotropic resolution and 230 mm × 230 mm field of view. For the T1-weighted scan, TR of 2.55 s, TE of 1.63 ms, inversion time (TI) of 1.15 s, flip angle of 8 degrees, and in-plane acceleration factor 2 were used. The T2-weighted scan was acquired with a TR of 3.2 s, a TE of 564 ms, and an in-plane acceleration factor of 2.

### Functional MRI Data Preprocessing

Results included in this manuscript come from preprocessing performed using *fMRIPrep* 23.1.3(Esteban et al., 2019; Esteban et al., 2023; RRID:SCR 016216), which is based on *Nipype* 1.8.6 (Gorgolewski et al., 2011; RRID:SCR 002502).

#### Anatomical data preprocessing

A total of 1 T1-weighted (T1w) image was found within the input BIDS dataset. The T1-weighted (T1w) image was corrected for intensity non-uniformity (INU) with N4BiasFieldCorrection (Tustison et al., 2010, distributed with ANTs (Avants et al., 2008, RRID:SCR 004757), and used as T1w-reference throughout the workflow. The T1w-reference was then skull-stripped with a *Nipype* implementation of the antsBrainExtraction.sh workflow (from ANTs), using OASIS30ANTs as the target template. Brain tissue segmentation of cerebrospinal fluid (CSF), white matter (WM), and gray matter (GM) was performed on the brain-extracted T1w using fast (FSL, RRID:SCR 002823, Zhang et al., 2001). Brain surfaces were reconstructed using recon-all (FreeSurfer 7.3.2, RRID:SCR 001847, Dale et al., 1999), and the brain mask estimated previously was refined with a custom variation of the method to reconcile ANTs-derived and FreeSurfer-derived segmentations of the cortical gray-matter of Mindboggle (RRID:SCR 002438, Klein et al., 2017). Volume-based spatial normalization to one standard space (MNI152NLin2009cAsym) was performed through nonlinear registration with antsRegistration (ANTs), using brain-extracted versions of both the T1w reference and the T1w template. The following template was selected for spatial normalization and accessed with *TemplateFlow* (23.0.0): *ICBM 152 Nonlinear Asymmetrical template version 2009c* (Fonov et al., 2009, RRID:SCR 008796; TemplateFlow ID: MNI152NLin2009cAsym).

#### Functional data preprocessing

A *B0* -nonuniformity map (or *fieldmap*) was estimated based on two (or more) echo-planar imaging (EPI) references with topup (Andersson et al., 2003; FSL). The estimated *fieldmap* was then aligned with rigid-registration to the target EPI (echo-planar imaging) reference run.

For each of the 6 BOLD runs found per subject (across all tasks and sessions), the following preprocessing was performed. First, a reference volume and its skull-stripped version were generated using a custom methodology of *fMRIPrep*. The BOLD reference was then co-registered to the T1w reference using bbregister (FreeSurfer) which implements boundary-based registration (Greve and Fischl, 2009). Co-registration was configured with six degrees of freedom. Head-motion parameters with respect to the BOLD reference (transformation matrices, and six corresponding rotation and translation parameters) are estimated before any spatiotemporal filtering using mcflirt (FSL, Jenkinson et al., 2002). BOLD runs were slice-time corrected to 0.52s (0.5 of the slice acquisition range 0s-1.04s) using 3dTshift from AFNI (Cox and Hyde, 1997, RRID:SCR 005927). The BOLD time-series were resampled onto the following surfaces (FreeSurfer reconstruction nomenclature): *fsaverage*. The BOLD time-series (including slice-timing correction when applied) were resampled onto their original, native space by applying a single, composite transform to correct for head-motion and susceptibility distortions. These resampled BOLD time-series will be referred to as *preprocessed BOLD in original space*, or just *preprocessed BOLD*. The BOLD time-series were resampled into standard space, generating a *preprocessed BOLD run in MNI152NLin2009cAsym space*. Several confounding time-series were calculated based on the *preprocessed BOLD* : framewise displacement (FD), DVARS, and three region-wise global signals.

FD was computed using two formulations following Power (absolute sum of relative motions, Power et al., 2014) and Jenkinson (relative root mean square displacement between affines, Jenkinson et al., 2002). FD and DVARS are calculated for each functional run, both using their implementations in *Nipype* (following the definitions by Power et al., 2014). The three global signals are extracted within the CSF, the WM, and the whole-brain masks. Additionally, a set of physiological regressors was extracted to allow for component-based noise correction (*CompCor*, Behzadi et al., 2007). Principal components are estimated after high-pass filtering the *preprocessed BOLD* time-series (using a discrete cosine filter with 128s cut-off) for the two *CompCor* variants: temporal (tCompCor) and anatomical (aCompCor). tCompCor components are then calculated from the top 2% variable voxels within the brain mask. For aCompCor, three probabilistic masks (CSF, WM, and combined CSF+WM) are generated in anatomical space. The implementation differs from that of Behzadi et al. (2007) in that instead of eroding the masks by 2 pixels on BOLD space, a mask of pixels that likely contains a volume fraction of GM is subtracted from the aCompCor masks. This mask is obtained by dilating a GM mask extracted from the FreeSurfer’s *aseg* segmentation, and it ensures components are not extracted from voxels containing a minimal fraction of GM. Finally, these masks are resampled into BOLD space and binarized by thresholding at 0.99 (as in the original implementation). Components are also calculated separately within the WM and CSF masks. For each CompCor decomposition, the *k* components with the largest singular values are retained, such that the retained components’ time series are sufficient to explain 50 percent of variance across the nuisance mask (CSF, WM, combined, or temporal). The remaining components are dropped from consideration. The head-motion estimates calculated in the correction step were also placed within the corresponding confounds file. The confound time series derived from head motion estimates and global signals were expanded with the inclusion of temporal derivatives and quadratic terms for each (Satterthwaite et al., 2013). Frames that exceeded a threshold of 0.5 mm FD or 1.5 standardized DVARS were annotated as motion outliers. Additional nuisance time-series are calculated by means of principal components analysis of the signal found within a thin band (*crown*) of voxels around the edge of the brain, as proposed by Patriat, Reynolds, and Birn (2017). All resamplings can be performed with *a single interpolation step* by composing all the pertinent transformations (i.e., head-motion transform matrices, susceptibility distortion correction when available, and co-registrations to anatomical and output spaces). Gridded (volumetric) resamplings were performed using antsApplyTransforms (ANTs), configured with Lanczos interpolation to minimize the smoothing effects of other kernels (Lanczos, 1964). Non-gridded (surface) resamplings were performed using mri vol2surf (FreeSurfer).

Many internal operations of *fMRIPrep* use *Nilearn* 0.10.1 [Abraham et al., 2014, RRID:SCR 001362], mostly within the functional processing workflow. For more details of the pipeline, see the section corresponding to workflows in *fMRIPrep*’s documentation (https://fmriprep.org/en/latest/workflows.html).

### Functional MRI Data Analysis

The SPM12 package was used for the GLM analysis on the fMRI data (Wellcome Department of Imaging Neuroscience, Institute of Neurology, London, UK). The fMRI data were slice-timing corrected and were applied with a high-pass filter of 180 seconds to remove low-frequency drifts potentially caused by physiological and physical noise. The fMRI data were corrected for motion, warped to the standard Montreal Neurological Institute (MNI) template, and smoothed with a Gaussian kernel (8mm FWHM) to mitigate individual anatomical differences.

We set up a general linear model (GLM) to perform voxel-based statistical modeling on the BOLD activity. The two blocks of the fMRI data were concatenated into one longer sequence, and there are, in total, seven event-related regressors with their associated parametric modulators in the GLM. The event-related regressors are modeled as a stick function with zero duration and are followed by the z-scored parametric modulators if they have any. The GLM specification is as follows:

1. Fixation Onset;
2. Stimulus Onset: a)MF Chosen value, b) MF Rejected value, c) MB Chosen value, d) MB Rejected value;
3. Left Response Onset;
4. Right Response Onset;
5. Planet-Pad Onset: a)State Prediction Error (Note: The event onset regressor is a combined regressor from the two events of interest, and the SPE signals at the planet and pad stage are combined into one parametric modulator);
6. Planet-Pad-Outcome Onset: a)MF Reward Prediction Error, b) MB Reward Prediction Error (Note: The event onset regressor is a combined regressor from the onsets of the three events, and the RPEs at the three stages are combined into one parametric modulator for the MF and MB systems, respectively);
7. Outcome Onset: a)Reward Magnitude.

The events of interest in the design matrix for each trial are stimulus onset, response onset, planet onset, pad onset, and outcome onset, which are set as stick functions at the corresponding event onset time. Since there are RPE signals from the MF and MB systems arising from three different stages on each trial, to increase the statistical power of capturing the RPE-specific variance in the BOLD signal, we combine the three event regressors into one chained regressor “planet-pad-outcome onset,” which essentially means that for each trial, there are three stick functions built into the combined regressor rather than one stick function per trial. The combined “planet-pad-outcome” regressor, which entails three stick functions, is parametrically modulated by the combined MF RPE and MB RPE regressors. Consequently, using the combined event regressor for the RPE signal would have three times more observation points than setting three separate RPE regressors at each stage (and it would not differentiate RPE encoding at different trial stages). Also, both MF RPE and MB RPE are simultaneously put as parametric modulators of the combined event regressor without orthogonalization, so that the identified signal ascribed to either MF or MB RPE regressor is beyond the shared variance and belongs to the regressor itself. The three-stage MF RPE and MB RPE signals are both z-scored across all trials, and we fitted the first-level GLM to each participant with the reward magnitude as a parametric modulator at the outcome onset stage to control for the confounding effect from the outcome magnitude. For data inclusion in this analysis, as we are interested in the RPE computations in RL, we excluded the Other Group and reported the results from analyzing the data of the three RL-related behavioral groups (i.e., the MB, Mixture, and MF Groups). For the SPE signals, as there are two stages where the SPEs would arise (i.e., planet and pad stage), the SPE regressor is also built into the GLM as a combined parametric regressor, similar to the RPE regressors.

To control for motion and non-neuronal fMRI signals, we include the following nuisance regressors: six rigid-body motion regressors (three translations and three rotations), framewise displacement (FD; quantification of the estimated bulk-head motion) (Power et al., 2012), the averaged signal within cerebrospinal fluid (CSF) mask, the average signal within the white matter mask, the average signal within both the cerebrospinal fluid and white matter mask, and the average signal within the whole brain mask and the derivative of the average signal within the whole brain mask. Besides, the scan volumes with an FD larger than the threshold of 0.77mm were set as regressors of motion spikes. The threshold of 0.77mm is determined by first calculating the FD threshold of 1.5 times the interquartile range plus the third quartile of each participant’s FD across all scans, then performing the calculation of 1.5 times the interquartile range plus the third quartile of the entire FD threshold distribution across all individuals.

#### Whole-brain voxel-based statistical analysis

To test the effects of decision value from the MB/MF system at the whole-brain voxel-wise level, we computed at the contrast level the betas of the MB/MF chosen value minus that of the MB/MF rejected value. The same procedure was carried out for each of the behavioral cluster groups (i.e., the MB, MF, Mixture, and Other group) to test the effects of the MB/MF decision value. A second-level GLM was run on the first-level betas to test the overall MB/MF decision value effect (one-sample t-test), and 4 separate second-level GLM analyses were run for each of the four behavioral groups to test such decision value effects in the corresponding group. To test where in the brain the decision value contrasts would vary across individuals (in three RL groups) with different behavioral MB (or MF) tendency, the model-derived variable, *wMF*, which is the averaged weight after adjustment for three reliability manipulations across all trials for each participant, was put as the parametric variable in the second-level GLM analysis. To determine whether there are any significantly different effects of the MB/MF decision value between the MB group and MF group, a whole-brain voxel-based two-sample t-test between the MF group and MB group was conducted on the first-level betas of the decision value contrasts.

For the investigation of the reward prediction error (RPE) signals in the brain, the second-level GLM was performed on the first-level betas of the RPE contrasts (i.e., MF RPE and MB RPE), firstly at the whole-brain voxel-based level. The statistical significance of the MF RPE and MB RPE signals was then examined in the corresponding regions of interest (i.e., caudate and ventral striatum) using small-volume correction. To examine the anatomical specificity of MF RPE and MB RPE within the striatum, the difference between the betas of the MF RPE and MB RPE contrasts was carried out through the one-sample t-test in second-level GLM. The RPE contrast testing at the sub-group level was conducted similarly to the decision value contrast testing through the second-level GLMs.

The representation of the state prediction error (SPE) signals in the brain was first examined through the whole-brain voxel-wise approach, similarly to the RPE signals. To detect where in the brain the SPE effects were modulated by an individual’s behavioral MB (or MF) tendency, we again ran a second-level GLM analysis with the parametric variable *wMF* included for each participant to perform whole-brain voxel-wise statistical testing.

#### ROI-based statistical analysis

For analyses of the MB and MF decision value signals in the vmPFC ROI, the first-level betas of the relevant contrasts were extracted from the vmPFC mask (Bartra et al., 2013; see the next section on Regions of Interest) for all participants. The existence of the decision value signals in each behavioral group was then tested through one-sample t-tests on the beta values within each group. For the statistical test on the relationship between the beta values and the behavioral MB (or MF) tendency entailed by the group identity, a regression model was run on the extracted beta values with an intercept and an ordinal group variable:

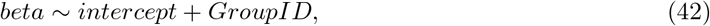

where “beta” denotes the beta values extracted from the ROI of the participants involved, and *GroupID* is an ordinal variable with “1” denoting the MB group, “2” denoting the Mixture group, and “3” denoting the MF group.

For the analysis of the relationship between the contrasts of interest in a given ROI and the behavioral MF tendency of each individual, Spearman correlations were conducted between the beta values extracted for the contrast in each ROI and the derived variable *wMF* from the computational model, which is a continuous measure of an individual’s behavioral MF (or MB) tendency.

### Regions of Interest and Small Volume Correction

The ROI analysis of MF and MB decision value within vmPFC is through using a vmPFC mask published from a meta-analysis on 206 fMRI studies of neural correlates of subjective value, where the mask is derived, across studies, from voxels that significantly showed a positive effect of values (Bartra et al., 2013).

For small volume correction conducted for the contrasts of MF RPE, the ROI mask of combined bilateral dorsal and ventral striatal regions was used as in the atlas from Pauli et al. (2018), which is composed of the bilateral caudate, putamen and nucleus accumbens.

For the ROI analysis of state prediction errors (SPE) in the bilateral intraparietal sulcus (IPS) and the bilateral dorsolateral prefrontal cortex (dlPFC), spherical ROIs were defined based upon the peak voxel coordinates of the average SPE signal found by Gläscher et al. (2010). Specifically, the ROI of bilateral IPS was defined as two 10 mm spheres centered at (-27, -54, 45) and (39, -54, 39), respectively. The ROI of bilateral dlPFC was also defined as two 10 mm spheres centered at (-45, 9, 33) and (45, 12, 30), respectively.

## Acknowledgements

We thank Julian Michael Tyszka from the Caltech Brain Imaging Center for assistance with the functional neuroimaging. We thank Emmily Hovhannisyan from UCLA for help in participant recruitment. We thank Rani Gera from Caltech for advice on data processing. This study was supported by a grant from the National Institute of Mental Health (R01MH121089) to J.P.O.D.

## Declaration of Interests

The authors declare no competing interests.

## Notes

### Competing Interest Statement

The authors have declared no competing interest.

### Summary of Updates

Small correction to the clustering membership and subsequent analyses; Reward prediction error results moved into the supplements; Figure reorganization.

