## Supplemental Materials for "Model-based and model-free valuation signals in the human brain vary markedly in their relationship to individual differences in human behavioral control"

### Supplementals

#### Figure S1: Performance and estimated MF weight of the 4 sub-groups

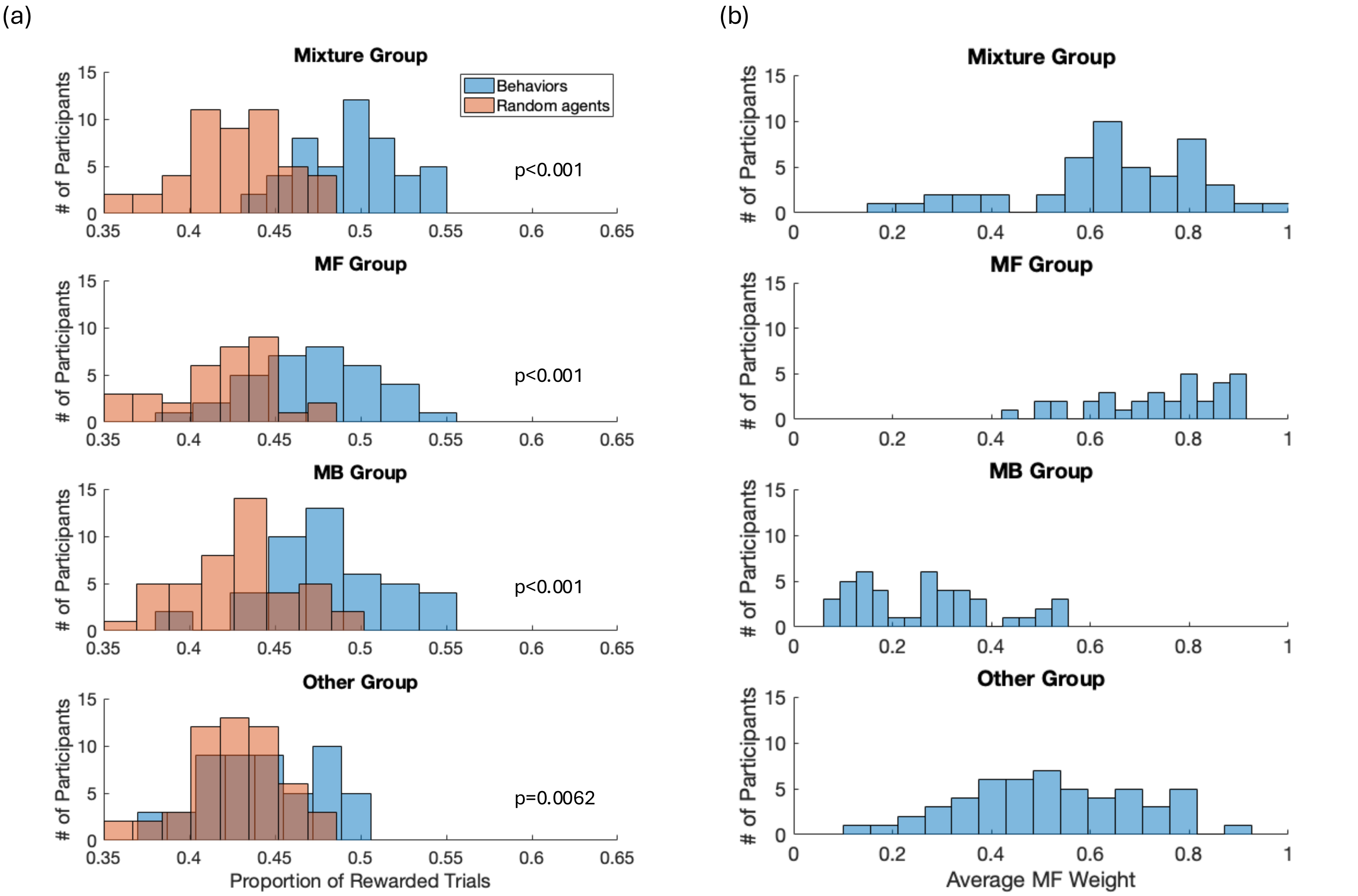


**Figure S1: Task performance and estimated MF weight (average wMF from the computational model) for the four sub-groups.** (a) The performance of the three RL groups (i.e., Mixture, MF and MB Groups) show significantly better task performance in reward rate compared to the random agents; the Other group showed slightly better-than-chance performance (blunted reward-driven behavior). The p-values show the two-sample test on difference between the sub-group behavior and the random agents' behaviors. (b) The average MF weight (wMF) distribution for each sub-group obtained from the clustering with Mixture, MF and MB Groups showing label-consistent wMF values, whereas the Other group showed the wMF centered around the prior value (0.5).

Table S1: Overall

| Model Parameter | Description | Mean ± SEM | Range |
|---|---|---|---|
| $\beta$ | Inverse temperature | $12.1662 \pm 6.7859$ | [0, ∞) |
| $\alpha_{MF}$ | MF value learning rate | $0.5649 \pm 0.0198$ | [0, 1] |
| $\alpha_{MB}$ | MB value learning rate | $0.5780 \pm 0.0221$ | [0, 1] |
| $wMF^{baseline}$ | Baseline MF weight parameter (before transformation) | $0.2586 \pm 0.1184$ | (-∞, ∞) |
| $a_{mbRPE}$ | Adjusted MF weight according to the reward contingency condition | $0.1399 \pm 0.0722$ | (-∞, ∞) |
| $a_{mfRPE}$ | Adjusted MF weight according to the reward magnitude condition | $-0.0032 \pm 0.0678$ | (-∞, ∞) |
| $a_{SPE}$ | Adjusted MF weight according to the state-transition uncertainty condition | $-0.0568 \pm 0.0641$ | (-∞, ∞) |
| $W_{MF}^{stay}$ | Stay bias for the previously selected option based on previous trial's outcome | $0.3726 \pm 0.0640$ | (-∞, ∞) |
| $W_{MF}^{switch}$ | Switch bias for the previously selected option based on previous trial's outcome | $0.1064 \pm 0.0447$ | (-∞, ∞) |
| $W_{MB}^{stay}$ | Stay bias for the previously selected option based on previous trial's outcome and transition | $0.4583 \pm 0.0605$ | (-∞, ∞) |
| $W_{MB}^{switch}$ | Switch bias for the previously selected option based on previous trial's outcome and transition | $0.1900 \pm 0.0412$ | (-∞, ∞) |
| $wMag$ | Weighting parameter of reward magnitude bias added to the terminal-state MB value | $0.8824 \pm 0.1342$ | [0, ∞) |

Table S2: Mixture Group

| Model Parameter | Description | Mean $\pm$ SEM | Range |
|---|---|---|---|
| $\beta$ | Inverse temperature* | 32.0812$\pm$25.2506 | [0, ∞) |
| $\alpha_{MF}$ | MF value learning rate | 0.5679$\pm$0.0319 | [0, 1] |
| $\alpha_{MB}$ | MB value learning rate | 0.6668$\pm$0.0369 | [0, 1] |
| $wMF^{baseline}$ | Baseline MF weight parameter (before transformation) | 0.9260$\pm$0.1853 | (-∞, ∞) |
| $a_{mbRPE}$ | Adjusted MF weight according to the reward contingency condition | 0.2149$\pm$0.1132 | (-∞, ∞) |
| $a_{mfRPE}$ | Adjusted MF weight according to the reward magnitude condition | -0.1074$\pm$0.1128 | (-∞, ∞) |
| $a_{SPE}$ | Adjusted MF weight according to the state-transition uncertainty condition | -0.0948$\pm$0.1224 | (-∞, ∞) |
| $W_{MF}^{stay}$ | Stay bias for the previously selected option based on previous trial's outcome | 0.6540$\pm$0.0862 | (-∞, ∞) |
| $W_{MF}^{switch}$ | Switch bias for the previously selected option based on previous trial's outcome | -0.0749$\pm$0.0374 | (-∞, ∞) |
| $W_{MB}^{stay}$ | Stay bias for the previously selected option based on previous trial's outcome and transition | 0.6677$\pm$0.1146 | (-∞, ∞) |
| $W_{MB}^{switch}$ | Switch bias for the previously selected option based on previous trial's outcome and transition | 0.0448$\pm$0.0526 | (-∞, ∞) |
| $wMag$ | Weighting parameter of reward magnitude bias added to the terminal-state MB value | 1.1067$\pm$0.2764 | [0, ∞) |

*Note: the Mean$\pm$ SEM inverse temperature is driven by a large observation in the sample ($\beta$>1000), without that participant, the Mean$\pm$ SEM becomes 6.8422 $\pm$ 0.7848.

Table S3: MF Group

| Model Parameter | Description | Mean ± SEM | Range |
|---|---|---|---|
| $\beta$ | Inverse temperature | 6.8966 ± 1.0485 | [0, ∞) |
| $\alpha_{MF}$ | MF value learning rate | 0.6193 ± 0.0533 | [0, 1] |
| $\alpha_{MB}$ | MB value learning rate | 0.4737 ± 0.0508 | [0, 1] |
| $wMF^{baseline}$ | Baseline MF weight parameter (before transformation) | 1.6448 ± 0.1817 | (-∞, ∞) |
| $a_{mbRPE}$ | Adjusted MF weight according to the reward contingency condition | -0.0562 ± 0.1739 | (-∞, ∞) |
| $a_{mfRPE}$ | Adjusted MF weight according to the reward magnitude condition | 0.2586 ± 0.1526 | (-∞, ∞) |
| $a_{SPE}$ | Adjusted MF weight according to the state-transition uncertainty condition | -0.1386 ± 0.1346 | (-∞, ∞) |
| $W_{MF}^{stay}$ | Stay bias for the previously selected option based on previous trial's outcome | 1.0091 ± 0.1526 | (-∞, ∞) |
| $W_{MF}^{switch}$ | Switch bias for the previously selected option based on previous trial's outcome | 0.2146 ± 0.1017 | (-∞, ∞) |
| $W_{MB}^{stay}$ | Stay bias for the previously selected option based on previous trial's outcome and transition | 0.0994 ± 0.1276 | (-∞, ∞) |
| $W_{MB}^{switch}$ | Switch bias for the previously selected option based on previous trial's outcome and transition | 0.1945 ± 0.1000 | (-∞, ∞) |
| $wMag$ | Weighting parameter of reward magnitude bias added to the terminal-state MB value | 0.9406 ± 0.4423 | [0, ∞) |

Table S4: MB Group

| Model Parameter | Description | Mean ± SEM | Range |
|---|---|---|---|
| $\beta$ | Inverse temperature | 5.6646±0.6106 | [0, ∞) |
| $\alpha_{MF}$ | MF value learning rate | 0.5176±0.0359 | [0, 1] |
| $\alpha_{MB}$ | MB value learning rate | 0.7166±0.0377 | [0, 1] |
| $wMF^{baseline}$ | Baseline MF weight parameter (before transformation) | -1.4282±0.1476 | (-∞, ∞) |
| $a_{mbRPE}$ | Adjusted MF weight according to the reward contingency condition | 0.1912±0.0997 | (-∞, ∞) |
| $a_{mfRPE}$ | Adjusted MF weight according to the reward magnitude condition | 0.3289±0.1045 | (-∞, ∞) |
| $a_{SPE}$ | Adjusted MF weight according to the state-transition uncertainty condition | -0.1978±0.0856 | (-∞, ∞) |
| $W_{MF}^{stay}$ | Stay bias for the previously selected option based on previous trial's outcome | 0.0793±0.1006 | (-∞, ∞) |
| $W_{MF}^{switch}$ | Switch bias for the previously selected option based on previous trial's outcome | 0.1160±0.1404 | (-∞, ∞) |
| $W_{MB}^{stay}$ | Stay bias for the previously selected option based on previous trial's outcome and transition | 1.0189±0.0931 | (-∞, ∞) |
| $W_{MB}^{switch}$ | Switch bias for the previously selected option based on previous trial's outcome and transition | 0.3286±0.0675 | (-∞, ∞) |
| $wMag$ | Weighting parameter of reward magnitude bias added to the terminal-state MB value | 0.4208± 0.1173 | [0, ∞) |

Table S5: Other Group

| Model Parameter | Description | Mean $\pm$ SEM | Range |
|---|---|---|---|
| $\beta$ | Inverse temperature | 2.9082$\pm$0.4034 | [0, ∞) |
| $\alpha_{MF}$ | MF value learning rate | 0.5664$\pm$0.0399 | [0, 1] |
| $\alpha_{MB}$ | MB value learning rate | 0.4495$\pm$0.0399 | [0, 1] |
| $wMF^{baseline}$ | Baseline MF weight parameter (before transformation) | 0.1652$\pm$0.1689 | (-∞, ∞) |
| $a_{mbRPE}$ | Adjusted MF weight according to the reward contingency condition | 0.1552$\pm$0.1731 | (-∞, ∞) |
| $a_{mfRPE}$ | Adjusted MF weight according to the reward magnitude condition | -0.3525$\pm$0.1436 | (-∞, ∞) |
| $a_{SPE}$ | Adjusted MF weight according to the state-transition uncertainty condition | 0.1470$\pm$0.1471 | (-∞, ∞) |
| $W_{MF}^{stay}$ | Stay bias for the previously selected option based on previous trial's outcome | -0.0470$\pm$0.1172 | (-∞, ∞) |
| $W_{MF}^{switch}$ | Switch bias for the previously selected option based on previous trial's outcome | 0.1931$\pm$0.0582 | (-∞, ∞) |
| $W_{MB}^{stay}$ | Stay bias for the previously selected option based on previous trial's outcome and transition | 0.0334$\pm$0.0901 | (-∞, ∞) |
| $W_{MB}^{switch}$ | Switch bias for the previously selected option based on previous trial's outcome and transition | 0.2036$\pm$0.0975 | (-∞, ∞) |
| $wMag$ | Weighting parameter of reward magnitude bias added to the terminal-state MB value | 1.0250$\pm$0.2278 | [0, ∞) |

Table S6: Trial-average MF weight overall and for each sub-group

| Overall/Sub-Group | Averaged MF weight (range: [0,1]) | SEM |
|---|---|---|
| Overall | 0.5322 | 0.0173 |
| Mixture | 0.6436 | 0.0259 |
| MF | 0.7332 | 0.0228 |
| MB | 0.2684 | 0.0210 |
| Other | 0.5215 | 0.0239 |

Note: The averaged MF weight is the mean of the trial-wise wMF variable across all trials used for model fitting. This trial-wise wMF was first calculated by adjusting the baseline wMF parameter through conditional weight changes (i.e., fitted parameters) and then fed into a sigmoid function to ensure it fell within the range of [0, 1] (for details of how the average MF weight is calculated, see “Computational Models” in the Methods section).

Table S7: MB Value (cluster-level FWE p<0.05)

| Peak coordinates (x, y, z) in mm | Hemisphere | Peak region | Cluster size | T-value/Z-value | P-value (cluster-level, FWE-corrected) |
|---|---|---|---|---|---|
| (-2, 56, -16) | Left | Ventromedial prefrontal cortex | 314 | 4.31/4.16 | P=0.002 |
| (-12, 58, 30) | Left | Frontal pole | 382 | 4.57/4.38 | P=0.001 |
| (10, 0, 44) | Right | Mid-cingulate cortex | 1592 | 5.13/4.88 | P<0.001 |
| (-14, -86, 26) | Left | Occipital pole / cuneal cortex | 2698 | 6.06/5.66 | P<0.001 |
| (20, -2 , -18) | Right | Amygdala | 404 | 5.95/5.57 | P<0.001 |
| (60, 2, 10) | Right | Precentral gyrus | 2288 | 6.13/5.71 | P<0.001 |
| (-50, -34, 20) | Left | Parietal operculum cortex | 2680 | 5.59/5.27 | P<0.001 |
| (-56, -6, -16) | Left | Middle temporal gyrus (anterior) | 606 | 5.10/4.85 | P<0.001 |
| (-48, 28, -4) | Left | Inferior frontal gyrus | 540 | 4.70/4.50 | P<0.001 |
| (-54, -6, 46) | Left | Precentral gyrus | 304 | 5.33/5.05 | P=0.003 |
| (-30, -28, -14) | Left | Hippocampus | 413 | 5.04/4.80 | P<0.001 |
| (-18, -44, 66) | Left | Postcentral gyrus | 384 | 4.62/4.43 | P=0.001 |

Table S8: MF Value (cluster-level FWE p<0.05)

| Peak coordinates (x, y, z) in mm | Hemisphere | Peak region | Cluster size | T-value/Z-value | P-value (cluster-level, FWE-corrected) |
|---|---|---|---|---|---|
| (-4, 62, 6) | Left | Frontal pole | 3985 | 6.67/6.16 | P<0.001 |
| (-62, -6, -6) | Left | Central opercular cortex | 4480 | 6.57/6.07 | P<0.001 |
| (20, -86, 38) | Right | Lateral occipital cortex / occipital pole | 6314 | 7.06/6.46 | P<0.001 |
| (62, 6, 0) | Right | Temporal pole | 2609 | 6.27/5.83 | P<0.001 |
| (50, -76, 16) | Right | Lateral occipital cortex (superior) | 171 | 4.49/4.32 | P=0.026 |
| (20, -44, 66) | Right | Postcentral gyrus | 279 | 4.68/4.49 | P=0.002 |
| (-22, -26, 58) | Left | Precentral gyrus | 274 | 4.22/4.07 | P=0.002 |
| (-52, -10, 40) | Left | Precentral gyrus | 146 | 3.89/3.77 | P=0.049 |

### Table S9:

#### The correlations between MB decision value and the degree of MB control

| Peak coordinates (x, y, z) in mm | Hemisphere | Peak region | Cluster size | T-value/Z-value | P-value (cluster-level, FWE-corrected) |
|---|---|---|---|---|---|
| (-6, 44, -12) | Left | Ventromedial prefrontal cortex | 1152 | 4.81/4.60 | P<0.001 |
| (-10, -56, 26) | Left | Precuneus cortex | 641 | 4.63/4.44 | P<0.001 |

**Table S9: The whole-brain voxel-wise results of significant correlations between MB decision value and the degree of MB control (-wMF).** The table lists the clusters that shows significant negative correlations between the representational strength of MB decision value and the degree of MF behavioral control (wMF), surviving cluster-level FWE-correction ($p<0.05$ with a cluster-forming threshold of $p<0.001$).

### Table S10: MB decision value in the MB group

| Peak coordinates (x, y, z) in mm | Hemisphere | Peak region | Cluster size | T-value/Z-value | P-value (cluster-level, FWE-corrected) |
|---|---|---|---|---|---|
| (-8, -56, 22) | Left | Precuneus cortex | 1916 | 6.94/5.65 | P<0.001 |
| (18, -2, -18) | Right | Amygdala | 232 | 6.72/5.52 | P=0.008 |
| (-6, 46, -14) | Left | Ventromedial prefrontal cortex | 2905 | 6.65/5.49 | P<0.001 |
| (-4, -18, 44) | Left | Mid-cingulate cortex | 331 | 6.51/5.40 | P=0.001 |
| (-58, -10, -12) | Left | Middle temporal gyrus (anterior) | 2657 | 6.14/5.17 | P<0.001 |
| (-48, 30, -2) | Left | Ventrolateral PFC | 855 | 5.48/4.74 | P<0.001 |
| (20, -86, 40) | Right | Lateral occipital cortex (superior) | 415 | 5.13/4.51 | P<0.001 |
| (66, -2, 8) | Right | Central opercular cortex | 1101 | 4.92/4.36 | P<0.001 |
| (-20, -2, -16) | Left | Amygdala | 190 | 4.66/4.17 | P=0.021 |
| (6, -6, 72) | Right | Supplementary motor cortex | 159 | 4.30/3.90 | P=0.044 |

**Table S10: MB decision value whole-brain voxel-wise SPM results in the MB group.** The table lists clusters correlating with the MB decision value in the MB group, surviving cluster-level FWE-correction ($p<0.05$ with a cluster-forming threshold of $p<0.001$).

### Table S11: MB decision value in the Mixture group

| Peak coordinates (x, y, z) in mm | Hemisphere | Peak region | Cluster size | T-value/Z-value | P-value (cluster-level, FWE-corrected) |
|---|---|---|---|---|---|
| (60, 0, 10) | Right | Precentral gyrus/ central opercular cortex | 185 | 4.67/4.21 | P=0.019 |
| (-48, -36, 18) | Left | Planum temporale/ parietal operculum cortex | 151 | 4.44/4.04 | P=0.044 |
| (-22, -64, 18) | Left | Supracalcarine cortex / precuneus cortex | 180 | 4.08/3.76 | P=0.021 |

**Table S11: MB decision value whole-brain voxel-wise SPM results in the Mixture group.** The table lists clusters correlating with the MB decision value in the Mixture group, surviving cluster-level FWE-correction ($p<0.05$ with a cluster-forming threshold of $p<0.001$).

**Note: There are no significant clusters surviving cluster-level FWE-correction ($p<0.05$ with a cluster-forming threshold of $p<0.001$) in either the MF or the Other group.**

### Table S12: MF decision value in the MB group

| Peak coordinates (x, y, z) in mm | Hemisphere | Peak region | Cluster size | T-value/Z-value | P-value (cluster-level, FWE-corrected) |
|---|---|---|---|---|---|
| (-4, 40, 0) | Left | Anterior cingulate cortex/ ventromedial prefrontal cortex | 562 | 4.94/4.37 | P<0.001 |
| (-8, -88, 40) | Left | Occipital pole | 304 | 4.56/4.10 | P=0.001 |
| (0, -10, 42) | Left & Right | Mid-cingulate cortex | 211 | 4.28/3.89 | P=0.010 |
| (-12, -42, -4) | Left | Lingual gyrus | 164 | 4.46/4.02 | P=0.032 |
| (6, -54, 2) | Right | Lingual gyrus | 175 | 5.38/4.68 | P=0.024 |
| (56, 14, -8) | Right | Temporal pole | 168 | 5.19/4.55 | P=0.029 |

**Table S12: MF decision value whole-brain voxel-wise SPM results in the MB group.** The table lists clusters correlating with the MF decision value in the MB group, surviving cluster-level FWE-correction ($p<0.05$ with a cluster-forming threshold of $p<0.001$).

### Table S13: MF decision value in the Mixture group

| Peak coordinates (x, y, z) in mm | Hemisphere | Peak region | Cluster size | T-value/Z-value | P-value (cluster-level, FWE-corrected) |
|---|---|---|---|---|---|
| (12, -90, 36) | Right | Occipital pole | 2258 | 6.76/5.62 | P<0.001 |
| (18, -62, -8) | Right | Lingual gyrus | 624 | 5.29/4.66 | P<0.001 |
| (-8, 56, 34) | Left | Frontal pole | 372 | 4.97/4.43 | P<0.001 |
| (-62, -34, 18) | Left | Planum temporale | 173 | 4.91/4.39 | P=0.029 |
| (64, -28, 22) | Right | Parietal operculum cortex | 249 | 4.90/4.38 | P=0.005 |
| (-46, -34, -4) | Left | Middle temporal gyrus (posterior) | 257 | 4.87/4.36 | P=0.004 |
| (-20, -68, -12) | Left | Occipital fusiform gyrus | 580 | 4.67/4.21 | P<0.001 |
| (-4, 52, -12) | Left | Ventromedial prefrontal cortex | 417 | 4.55/4.12 | P<0.001 |
| (48, -78, 12) | Right | Lateral occipital cortex | 212 | 4.18/3.83 | P=0.012 |

**Table S13: MF decision value whole-brain voxel-wise SPM results in the Mixture group.** The table lists clusters correlating with the MF decision value in the Mixture group, surviving cluster-level FWE-correction (p<0.05 with a cluster-forming threshold of p<0.001).

### Table S14: MF decision value in the MF group

| Peak coordinates (x, y, z) in mm | Hemisphere | Peak region | Cluster size | T-value/Z-value | P-value (cluster-level, FWE-corrected) |
|---|---|---|---|---|---|
| (-66, -16, 2) | Left | Superior temporal gyrus (posterior) | 645 | 5.43/4.55 | P<0.001 |
| (62, 4, -6) | Right | Superior temporal gyrus (anterior) | 182 | 5.90/4.84 | P=0.018 |
| (-6, 64, -2) | Left | Medial prefrontal cortex/frontal pole | 641 | 5.49/4.60 | P<0.001 |

**Table S14: MF decision value whole-brain voxel-wise SPM results in the MF group.** The table lists clusters correlating with the MF decision value in the MF group, surviving cluster-level FWE-correction ($p<0.05$ with a cluster-forming threshold of $p<0.001$).

### Table S15: MF decision value in the Other group

| Peak coordinates (x, y, z) in mm | Hemisphere | Peak region | Cluster size | T-value/Z-value | P-value (cluster-level, FWE-corrected) |
|---|---|---|---|---|---|
| (-6, 50, -4) | Left | Ventromedial prefrontal cortex / medial prefrontal cortex | 833 | 5.36/4.76 | P<0.001 |
| (-10, -46, 38) | Left | Posterior cingulate cortex/precuneus cortex | 202 | 4.59/4.18 | P=0.017 |

**Table S15: MF decision value whole-brain voxel-wise SPM results in the Other group.** The table lists clusters correlating with the MF decision value in the Other group, surviving cluster-level FWE-correction (p<0.05 with a cluster-forming threshold of p<0.001).

### Table S16: MB decision value, MB group > MF group

| Peak coordinates (x, y, z) in mm | Hemisphere | Peak region | Cluster size | T-value/Z-value | P-value (cluster-level, FWE-corrected) |
|---|---|---|---|---|---|
| (-8, 46, -16) | Left | Ventromedial prefrontal cortex | 455 | 4.70/4.39 | P<0.001 |
| (2, -62, 18) | Right | Precuneus cortex | 315 | 3.92/3.73 | P=0.002 |

**Table S16: MB decision value whole-brain voxel-wise SPM results for the contrast MB group > MF group.** The table lists clusters that show the stronger MB decision value encoding in the MB group than in the MF group, surviving cluster-level FWE-correction (p<0.05 with a cluster-forming threshold of p<0.001).

**Note: In terms of the MF decision value SPM results compared across the MF group and MB group, there are no significant clusters surviving cluster-level FWE-correction in vmPFC (p<0.05 with a cluster-forming threshold of p<0.001).**

### Table S17: State prediction errors across the whole brain

| Peak coordinates (x, y, z) in mm | Hemisphere | Peak region | Cluster size | T-value/Z-value | P-value (cluster-level, FWE-corrected) |
|---|---|---|---|---|---|
| (-30, -54, 42) | Left | Superior parietal lobule | 15797 | 9.03/Inf | P<0.001 |
| (44, 10, 32) | Right | Precentral gyrus/ middle frontal gyrus | 2657 | 6.90/6.34 | P<0.001 |
| (-42, 6, 30) | Left | Precentral gyrus/ dorsolateral prefrontal cortex | 1380 | 6.48/6.00 | P<0.001 |
| (-32, 14, 56) | Left | Middle frontal gyrus | 894 | 5.64/5.31 | P<0.001 |

**Table S17: State prediction errors whole-brain voxel-wise SPM results.** The table lists clusters correlating with the state prediction errors, surviving cluster-level FWE-correction (p<0.05 with a cluster-forming threshold of p<0.001).

### Table S18: State prediction errors in the MB group

| Peak coordinates (x, y, z) in mm | Hemisphere | Peak region | Cluster size | T-value/Z-value | P-value (cluster-level, FWE-corrected) |
|---|---|---|---|---|---|
| (-42, 4, 34) | Left | Dorsolateral PFC | 328 | 5.65/4.86 | P=0.001 |
| (46, 10, 34) | Right | Dorsolateral PFC | 888 | 5.38/4.68 | P<0.001 |
| (-30, -52, 42) | Left | Intraparietal sulcus | 6213 | 6.98/5.68 | P<0.001 |
| (6, 26, 44) | Right | Paracingulate gyrus | 216 | 4.63/4.15 | P=0.010 |
| (40, -58, -10) | Right | Temporal occipital fusiform cortex | 760 | 7.37/5.90 | P<0.001 |
| (34, 2, 62) | Right | Middle frontal gyrus | 812 | 6.58/5.44 | P<0.001 |
| (-28, -2, 56) | Left | Middle frontal gyrus | 637 | 6.20/5.21 | P<0.001 |
| (30, 24, 6) | Right | Insular cortex | 153 | 5.11/4.49 | P=0.045 |
| (-46, 28, 34) | Left | Middle frontal gyrus | 243 | 4.90/4.34 | P=0.005 |
| (-32, -80, -8) | Left | Occipital fusiform gyrus | 225 | 4.65/4.16 | P=0.008 |

**Table S18: State prediction errors whole-brain voxel-wise SPM results in the MB group.** The table lists clusters correlating with the state prediction errors in the MB group, surviving cluster-level FWE-correction ($p<0.05$ with a cluster-forming threshold of $p<0.001$).

### Table S19: State prediction errors in the Mixture group

| Peak coordinates (x, y, z) in mm | Hemisphere | Peak region | Cluster size | T-value/Z-value | P-value (cluster-level, FWE-corrected) |
|---|---|---|---|---|---|
| (30, -50, -16) | Right | Temporal occipital fusiform cortex | 1689 | 7.43/6.01 | P<0.001 |
| (-36, -74, 44) | Left | Lateral occipital cortex (superior) | 2312 | 7.28/5.92 | P<0.001 |
| (-40, -70, -14) | Left | Occipital fusiform gyrus | 648 | 7.05/5.79 | P<0.001 |
| (38, -78, 36) | Right | Lateral occipital cortex (superior) | 3557 | 6.97/5.75 | P<0.001 |
| (30, 12, 48) | Right | Middle frontal gyrus | 478 | 5.62/4.89 | P<0.001 |
| (-54, 12 ,44) | Left | Middle frontal gyrus/dorsolateral prefrontal cortex | 683 | 5.27/4.65 | P<0.001 |
| (-28, 12, 56) | Left | Middle frontal gyrus | 277 | 5.10/4.53 | P=0.002 |
| (50, 20, 36) | Right | Middle frontal gyrus/dorsolateral prefrontal cortex | 204 | 4.17/3.82 | P=0.012 |

**Table S19: State prediction errors whole-brain voxel-wise SPM results in the Mixture group.** The table lists clusters correlating with the state prediction errors in the Mixture group, surviving cluster-level FWE-correction (p<0.05 with a cluster-forming threshold of p<0.001).

### Table S20: State prediction errors in the MF group

| Peak coordinates (x, y, z) in mm | Hemisphere | Peak region | Cluster size | T-value/Z-value | P-value (cluster-level, FWE-corrected) |
|---|---|---|---|---|---|
| (24, -60, 52) | Right | Lateral occipital cortex (superior) | 295 | 6.40/5.12 | P=0.001 |
| (44, -78, 24) | Right | Lateral occipital cortex (superior) | 375 | 5.01/4.29 | P<0.001 |
| (-28, -68, 52) | Left | Lateral occipital cortex (superior) | 156 | 4.65/4.05 | P=0.036 |
| (-36, -58, -16) | Left | Temporal occipital fusiform cortex | 178 | 4.25/3.76 | P=0.020 |

**Table S20: State prediction errors whole-brain voxel-wise SPM results in the MF group.** The table lists clusters correlating with the state prediction errors in the MF group, surviving cluster-level FWE-correction (p<0.05 with a cluster-forming threshold of p<0.001).

### Table S21: State prediction errors in the Other group

| Peak coordinates (x, y, z) in mm | Hemisphere | Peak region | Cluster size | T-value/Z-value | P-value (cluster-level, FWE-corrected) |
|---|---|---|---|---|---|
| (32, -54, -8) | Right | Temporal occipital fusiform cortex | 453 | 6.32/5.42 | P<0.001 |

**Table S21: State prediction errors whole-brain voxel-wise SPM results in the Other group.** The table lists clusters correlating with the state prediction errors in the Other group, surviving cluster-level FWE-correction (p<0.05 with a cluster-forming threshold of p<0.001).

Table S22:

The correlations between state prediction errors and the degree of MB control

| Peak coordinates (x, y, z) in mm | Hemisphere | Peak region | Cluster size | T-value/Z-value | P-value (cluster-level, FWE-corrected) |
|---|---|---|---|---|---|
| (-38, -52, 58) | Left | Intraparietal sulcus /superior parietal lobule | 230 | 4.34/4.18 | P=0.013 |
| (-22, 4, 52) | Left | Superior frontal gyrus | 200 | 3.78/3.67 | P=0.025 |
| (30, 0, 54) | Right | Superior frontal gyrus | 228 | 4.40/4.24 | P=0.014 |

**Table S22: The whole-brain voxel-wise results of significant correlations between state prediction errors and the degree of MB control (-wMF).** The table lists the clusters that show significant negative correlations between the representational strength of state prediction errors and the degree of MF behavioral control (wMF), surviving cluster-level FWE-correction ($p<0.05$ with a cluster-forming threshold of $p<0.001$).

### Figure S2: Reward prediction errors in the striatum (signals at the combined 3 stages)

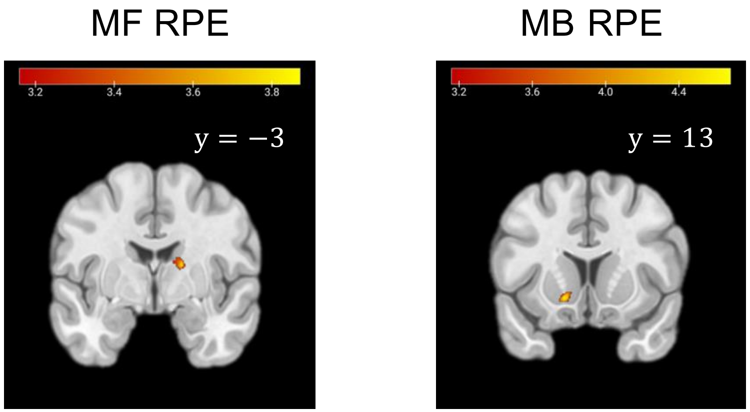


**Figure S2: The clusters of MF RPE and MB RPE contrasts in the striatum.** Group-level T-maps of MF RPE and MB RPE contrasts, with signals at the 1st ,2nd and 3rd (outcome) stages combined, in the caudate and ventral striatum ROIs, respectively. The clusters shown are obtained with a cluster-forming threshold of p<0.001 (uncorrected).

### Figure S3: Chosen & Rejected MB & MF Values

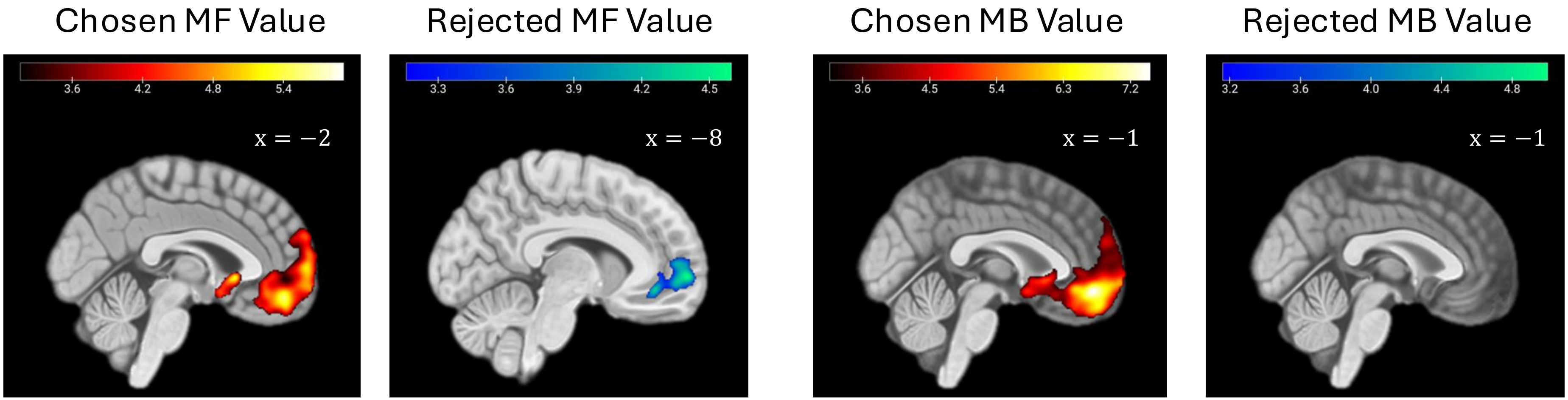


**Figure S3: The effects of chosen and rejected MB & MF values in the prefrontal cortex (the sub-components of the decision value contrasts).** Group-level T-maps in the prefrontal cortex survive cluster-level FWE correction ($p<0.05$ with a cluster-forming threshold of $p<0.001$). The chosen and rejected MF value as the sub-component of MF decision value are both significantly encoded in the prefrontal cortex; in contrast, for the sub-components of MB decision value, only the chosen but not the rejected MB value is significantly encoded.

### Table S23: Chosen and Rejected MB & MF value in Figure S2

| Contrasts | Peak coordinates (x, y, z) in mm | Hemisphere | Peak region | Cluster size | T-value/Z-value | P-value (cluster-level, FWE-corrected) |
|---|---|---|---|---|---|---|
| Chosen MF Value | (2, 16, -2) | Right | Subcallosal cortex | 4487 | 5.99/5.60 | P<0.001 |
| Rejected MF Value | (-8, 60, 0) | Left | Prefrontal cortex/ Frontal pole | 942 | 4.71/4.51 | P<0.001 |
| Chosen MB Value | (0, 48, -10) | Left & Right | Ventromedial prefrontal cortex | 13514 | 7.55/6.84 | P<0.001 |
| Rejected MB Value | / | / | / | / | / | / |

**Table S23: Cluster and peak voxel information of the chosen and rejected MB & MF value contrasts in Figure S3.**

### Figure S4: Model identifications in the current 2-step task

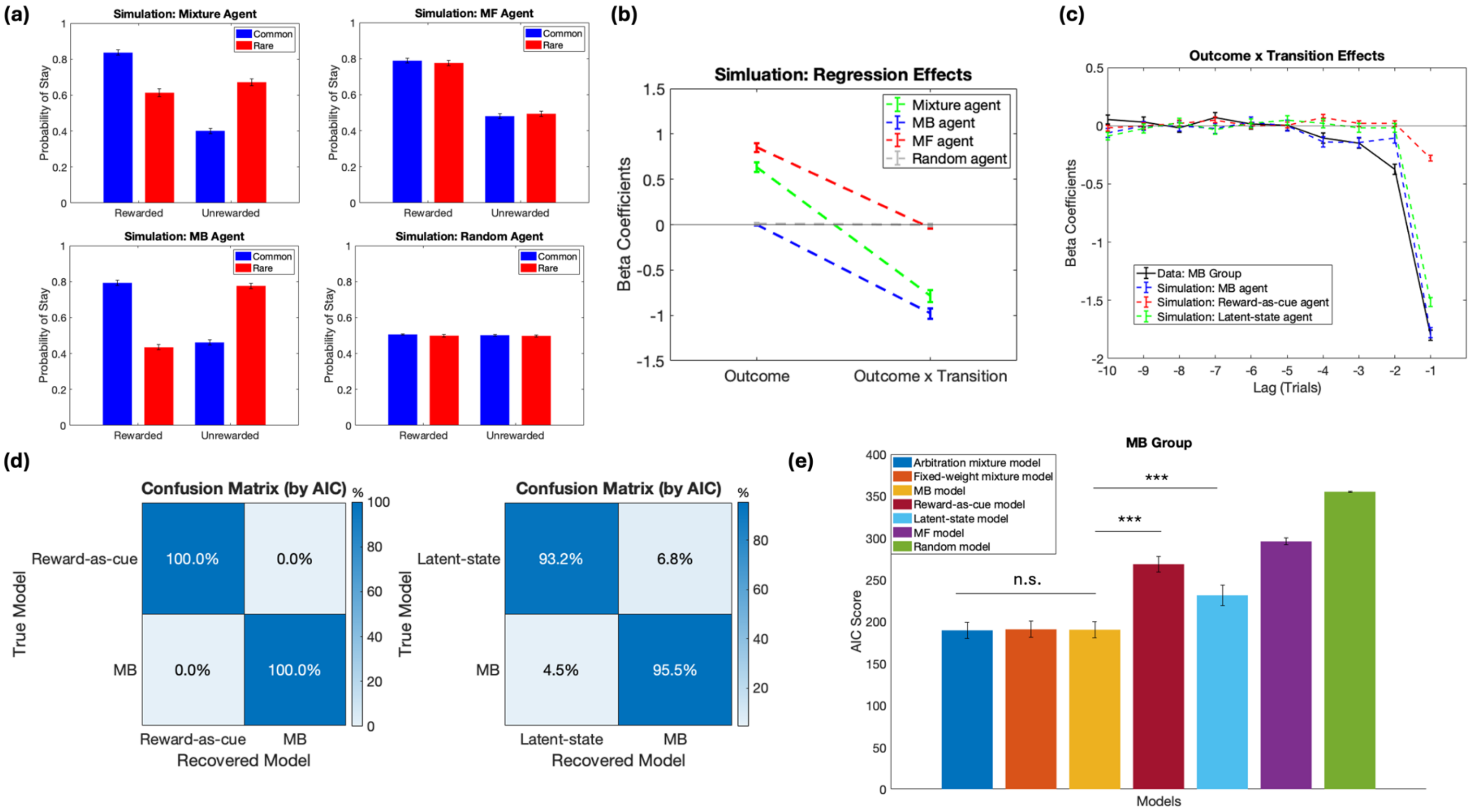


**Figure S4: Simulations of different agents in the current 2-step task and dissociation of the MB RL model from alternative algorithms in the MB group, addressing potential confounds brought up by Akam et al. (2015).** (a) The simulated choices of four agents when navigating the task, with each showing distinct stay choice patterns as a function of the previous trial's outcome and transition type. (b): The regression effects of outcome and outcome-transition interaction shown by the simulated agents, with each agent showing distinct and predicted composition of the regression effects in the simulated data. (c): The outcome-transition effects of last 10 trials on the current choice from both the actual data and the simulated agents, besides the MB agent, that could all show a preceding-trial outcome-transition effect on the current choice. (d):The identification sensitivity of the MB strategy, the alternative reward-as-cue model, and the latent-state model based on AIC score: the identification of MB strategy out of the aforementioned alternative models based on AIC scores has high accuracy. (e): Model comparison based upon AIC scores within the MB Group, where the Arbitration/Fixed-weight mixture model and the MB model are equally best at explaining the actual behaviors, better than the reward-as-cue or the latent-state model (see pages 29-30 and Akam et al. (2015) for descriptions of the reward-as-cue model and the latent-state model).

### Simulation and model recovery procedure

**Simulation**: all simulations were conducted by 1) using the set of fitted parameters to real behaviors to generate the corresponding population data for each agent (with the fitted parameters being the appropriate values for real behaviors), and then 2) regression analysis and model fitting were run on the generated data.

**Model recovery:** for two pairs of model identification: 1) the MB model vs. the reward-as-cue model, and 2) the MB model vs. the latent-state model, datasets of the true model were first generated as in the simulation procedure, then both the true model and the alternative were fitted to the generated datasets. Using the AIC measure, the proportion of participants generated from the true model, having a better fit from the true or the alternative model, was calculated to create the confusion matrix for each pair of model identifications.

### Reward-as-cue model

The reward-as-cue model treats all combinations of outcome (reward or no-reward) and second-stage state (red or green planet) in the previous trial as four separate states, and learns the two actions' Q-values in the current trial as a function of these four states using the Q-learning algorithm as described for the MF model.

$$RPE_t = O_t - Q_{t-1}(s_t, a_t);$$
$$Q_t(s_t, a_t) = Q_{t-1}(s_t, a_t) + \alpha \cdot RPE_t,$$

where $O_t$ denotes the current trial's magnitude outcome, $s_t$ denotes the one of the four states encountered in the previous trial ($s_t \in \{red - reward, green - reward, red - no - reward, green - no - reward\}$), and $a_t$ denotes the two options to choose in the current trial ($a_t \in \{choosing - yellow, choosing - blue\}$).

The action selection process is implemented by modeling the action probability through a softmax function on the corresponding Q-values for the relevant states among the four states:

$$P_t(s_t, a_t) = \frac{e^{\beta \cdot Q_t(s_t, a_t)}}{e^{\beta \cdot Q_t(s_t, a_t)} + e^{\beta \cdot Q_t(s_t, a'_t)}}, a_t \neq a'_t;$$

where $a_t, a'_t \in \{choosing - yellow, choosing - blue\}$.

Overall, the model has two free parameters: the softmax inverse temperature $\beta$ and the model-free learning rate α.

### Latent-state model

The latent-state model assumes there are two states of the world, $z_t$=1: the red planet is more likely to deliver rewards than the green planet; $z_t$=2: the green planet is more likely to deliver rewards than the red planet. It infers the probability that each of the two states is true by performing Bayesian updates to obtain the posterior probability based on events in each trial.

For each trial, assuming there is a probability $a$ that the current state would stay the same as in the previous trial and a probability $(1-a)$ that the current state would switch to the other state, the prior belief of being in one state is:

$$b_t(j) = \sum_{i=1}^{2} A_{ij}\alpha_{t-1}(i), \qquad A = \begin{bmatrix} a & 1-a \\ 1-a & a \end{bmatrix},$$

The $\alpha_t$ is the posterior belief for trial t, and it is updated through Bayes rule for each trial event:

$$\alpha_t(j) = \frac{b_t(j)B_t(j)}{\sum_{k=1}^{2} b_t(k)B_t(k)},$$

Where $B_t(j)$ denotes the Bernoulli likelihood:

$$B_t(1)\text{=P}(y_t|z_t = 1) = (q_t^1)^{y_t}\,(1-q_t^1)^{1-y_t}\ ; B_t(2)\text{=P}(y_t|z_t = 2) = (q_t^2)^{y_t}\,(1-q_t^2)^{1-y_t};$$

specifically, $y_t$ denotes the binary outcome, and the reward probability set $q_t^1$ and $q_t^2$ have the following form:

$$q_t^1 = \begin{cases} p_H, & \textit{if observing red planet} \\ p_L, & \textit{if observing green planet} \end{cases};$$

$$q_t^2 = \begin{cases} p_L, & \textit{if observing red planet} \\ p_H, & \textit{if observing green planet} \end{cases};$$

Where $p_L = 1 - p_H$, and $p_H, p_L$ are constrained in ranges of (0.5,1) and (0,0.5), respectively.

For choice selection, with the posterior probability of the two states $\alpha_t$, the action often leading to the high-reward planet in the more probable state is chosen with a probability of $1-\varepsilon$; and the action often leading to the high-reward planet in the less probable state is chosen with a probability of $\varepsilon$, with $\varepsilon$ in the range of (0,1).

Then the log-likelihood of actual choices are maximized for model fitting. In total, there are three free parameters: the state-stickiness $a$, the reward probability of the more rewarding planet $p_H$, and the choice randomness $\varepsilon$.
